# Same Movements, Different Skills: How Context Shapes Learning in Humans and Neural Networks

**DOI:** 10.64898/2026.11.08.764653

**Authors:** Mohan W. Gupta, Jordan A. Taylor

## Abstract

What determines whether experiences are integrated into shared representations or split? Motor skill learning offers a revealing test: when two tasks require an identical sequence of movements, both intuition and theory suggest that practice in one should generalize to the other. Yet we find that changing context can make identical movements different skills. In simple recurrent networks trained on the same sequence across contexts, splitting was greatest when strong contextual inputs coincided with a high initial weight scale; neither alone sufficed. Greater representational separation coincided with separate context-specific learning curves. Humans practicing the same finger sequence while alternating between two performance-irrelevant contexts likewise exhibited context-specific curves, rather than the single continuous trajectory observed when practicing in a single context. State-space modeling favored split learning states in humans and very weakly coupled states in the most context-dependent networks. These findings have broader implications for understanding how learning generalizes across contexts.

## Introduction

Did you know that *Twinkle, Twinkle Little Star* and *Baa Baa Black Sheep* have the same melody? If not, you are in good company. Prior work has found that we represent songs (melody and lyrics) in a bounded fashion (Crowder et al., 1990; Serafine et al., 1984, 1986). Their shared melody invites generalization, yet different lyrics can carve out separate memories. This highlights a tension in learning: when are experiences lumped together to support generalization, and when are they split to preserve specificity? Does this tension extend to the movements used to play a song? Would a budding pianist who has mastered *Twinkle, Twinkle Little Star* generalize that learning when playing *Baa Baa Black Sheep*, or would they have to start anew?

Motor sequence learning is often framed as acquiring a task-specific skill that becomes more fluent with practice and generalizes across changes in cueing, framing, or stimulus format (Keele et al., 1995; Willingham et al., 2000; Hikosaka et al., 2002; Ungerleider, 2002; Doyon et al., 2009; Dayan & Cohen, 2011; Abrahamse et al., 2013). Yet, real-world skills are rarely learned sans context. The same motor sequence is frequently learned in distinct contexts that can shape what is retrieved and expressed and, in some cases, changes what generalizes (Wright & Shea, 1991; Ruitenberg, Abrahamse, et al., 2012; L. Schmidt et al., 2012; Howard et al., 2013; Česonis & Franklin, 2022). This raises a basic question for motor learning: under what conditions does a learner lump together experiences across contexts for the same motor sequence and when do they treat them as separate skills that are split from each other?

We frame this question as a trade-off between lumped and split representations (Figure 1). In a lumped solution, practice in either context updates one shared representation. The sequence need only be learned once, so performance should improve smoothly across context changes, producing little or no transition cost (Figure 1A–B). In a split solution, the same sequence is represented separately across contexts (Figure 1C–D). Learning in one context does not fully update the representation used in the other. Switching contexts should therefore interrupt the learning trajectory and produce transition costs. The learner effectively maintains context-specific versions of the same skill.

**Figure 1.**
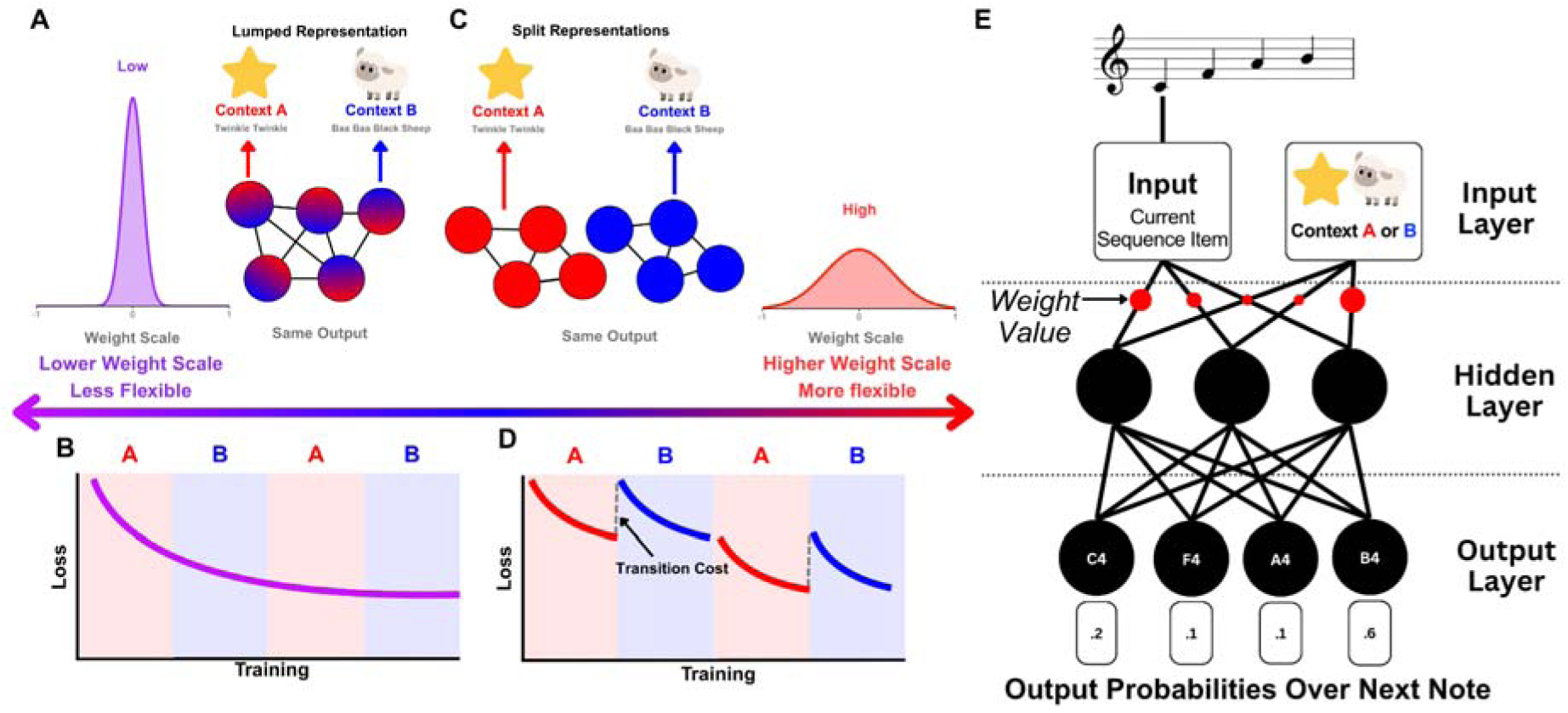
**(A)** With lower initial weight scale, networks are hypothesized to be less flexible and to favor a lumped representation, in which Contexts A and B rely on overlapping hidden-state representations for the same sequence. **(B)** A lumped representation predicts continuous improvement across context changes because learning in either context updates the same underlying representation. **(C)** With higher initial weight scale, networks are hypothesized to be more flexible and able to form split, context-specific representations while producing the same sequence output. **(D)** Split representations predict context-specific learning trajectories and transition costs when the context changes because learning does not fully generalize across representations. **(E)** SRN architecture. At each time step, the network receives the current sequence item and a context cue, updates its recurrent hidden state, and outputs probabilitie over the next sequence item. Initial weight scale was manipulated to vary the network’s representational flexibility.

This trade-off may help explain mixed evidence for context-dependent sequence memory. Effects vary with whether contextual features are incidental or task-relevant and whether performance is stimulus-driven or memory-based (Wright & Shea, 1991; Bapi et al., 2000; Ruitenberg, Abrahamse, et al., 2012; Sobierajewicz et al., 2017). These conditions may determine whether contextual features become incorporated into the learned skill or remain background information that can be ignored (Keele et al., 1995). When contextual cues provide little predictive structure, practice across contexts should be more likely to update a common sequence representation. When those cues are more predictive, they may instead become part of what is learned, providing a basis for splitting the same skill across contexts.

However, a strong context signal alone may not be sufficient to produce splitting. The learning system must also be able to reshape its internal representations around that signal. This idea is consistent with computational accounts in which learners infer whether experiences belong to the same underlying state or to distinct latent causes (Gershman et al., 2010; Gershman & Niv, 2010), as well as motor-control models in which separate internal models are selected and updated according to contextual or responsibility signals (Haruno et al., 2001; Heald et al., 2021). A system constrained by its existing representations may continue to update a shared skill despite contextual differences. A more flexible system could instead use context to partition experience. Thus, context-dependent splitting may depend both on the importance assigned to contextual information and on whether the learning system can reorganize around it.

Simple recurrent networks (SRNs; Elman networks) provide a minimal testbed for this hypothesis because their internal representations are learned, manipulable, and directly observable. Indeed, sequence learning was one of the first applications of SRNs, with early work showing that these networks can learn temporal and statistical structure by using recurrent hidden states to carry information forward in time (Servan-Schreiber et al., 1988; Elman, 1990; Cleeremans & McClelland, 1991). We trained SRNs to predict the next item in a fixed sequence, analogous to predicting the next note in a melody (Figure 1E). At each step, a network received the current sequence item and a context cue and used its recurrent hidden state to make the next prediction. Critically, the sequence and prediction target were identical across contexts, so context was unnecessary for successful performance. The network could therefore represent the sequence similarly across contexts, producing a lumped solution, or develop separate context-specific hidden-state trajectories, producing a split solution.

We leveraged this class of SRNs to identify the conditions that determine which of these representational solutions emerges. Prior work suggests that initial weight scale—the variance of a network’s starting weights—can change how much its internal representations reorganize during learning (Chizat et al., 2019; Dominé et al., 2025; Holton et al., 2026; Woodworth et al., 2020). We therefore used weight scale as an operational manipulation of network flexibility. In Low-Scale networks, network flexibility was expected to remain more constrained by the network’s initial representational structure, favoring lumping. In High-Scale networks, the network could more substantially reshape its hidden-state geometry, allowing context to organize separate representations. But greater flexibility alone might not produce splitting without a sufficiently strong context signal. We therefore manipulated both initial weight scale and context-signal strength, predicting the strongest splitting when both were high.

We first swept initial weight scale and context-signal strength to identify when splitting emerged in SRNs, examining both transition costs and internal representations. We then tested the resulting predictions in human motor sequence learning. Participants practiced the same finger sequence either within a single context or while alternating between two contexts unnecessary for successful performance. Finally, we fit lumped, cross-talk, and split state-space models to human response times and SRN loss to compare their learning dynamics within a common framework. Together, these approaches test when practice across contexts strengthens a shared skill and when identical movements become context-specific skills.

## Results

### SRNs reveal behavioral signatures of context-dependent splitting

Here, we sought to determine when SRNs lump or split the same sequence across contexts. We trained SRNs to predict the next item in a fixed eight-item sequence while alternating between Contexts A and B. Each network received the current sequence item and a context cue, although the sequence and prediction targets were identical across contexts. We predicted that networks forming a lumped representation would generalize across context switches with little or no transition cost (Figure 1A–B), whereas networks forming split representations would show larger transition costs (Figure 1C–D). We further hypothesized that splitting would depend on two factors: initial weight scale, which we used to manipulate network flexibility, and context strength. Specifically, we predicted that substantial splitting would emerge when high initial weight scale was paired with a strong context signal. We first swept both parameters across a broad range to determine where transition costs emerged.

Transition costs increased nonlinearly with weight scale, which affected both weight values and context signal values (Figure 2A). A mixed-effects model with random intercepts for network seed showed that a quadratic model fit substantially better than a linear model, χ^2^(1) = 58.32, *p* < .001. Transition costs increased with log weight scale, β = 0.350, *t*(139.2) = 10.55, *p* < .001, with additional positive curvature, β = 0.406, *t*(139.3) = 8.47, *p* < .001. Thus, transition costs remained small across much of the low-scale range before rising sharply at larger weight scales.

**Figure 2.**
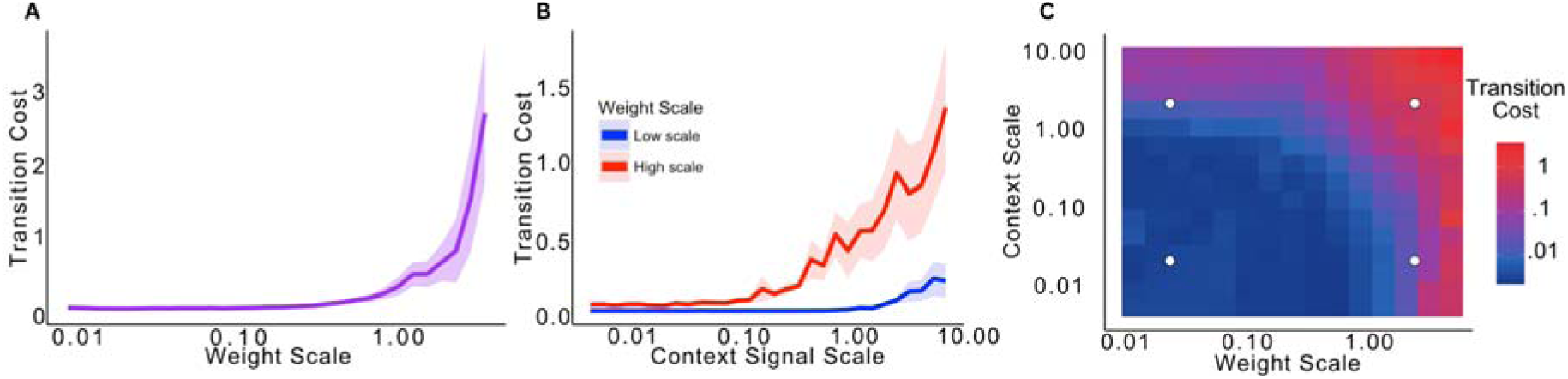
**(A)** Transition cost as a function of initial weight scale. Costs remained small across much of the low-scale regime before increasing sharply at larger weight scales. The line shows the mean across network runs and the shaded region shows ±1 SE. **(B)** Transition cost as a function of context-signal scale for Low-Scale and High-Scale networks. Increasing context strength produced little disruption in Low-Scale networks but increasingly large transition costs in High-Scale networks. Lines show mean across network runs and shaded regions show ±1 SE. **(C)** Joint sweep of initial weight scale and context-signal strength. Each tile represents the mean transition cost across network runs for a given parameter combination. Colors are mapped logarithmically from blue (lowest transition cost) to red (highest transition cost), with intermediate colors indicating increasing costs. Larger transition costs were concentrated where both weight scale and context-signal strength were high. White circles mark the four parameter combinations selected for the subsequent 2 × 2 design. Both axes are logarithmic.

We next swept context-signal strength separately in Low- and High-Scale networks. Increasing context strength had only a modest effect when weight scale was low, but produced much larger transition costs when weight scale was high (Figure 2B). A quadratic mixed-effects model again fit better than a linear model, χ^2^(2) = 70.50, *p* < .001. Critically, context strength interacted with weight scale, β = 0.282, *t*(295) = 10.55, *p* < .001, and this interaction became stronger at larger context values, as indicated by the weight-scale × quadratic context term, β = 0.150, *t*(295) = 4.81, *p* < .001.

A joint sweep of both parameters showed the same pattern across the full parameter space (Figure 2C): transition costs increased with both weight scale, β = 0.238, *t*(443) = 14.10, *p* < .001, and context strength, β = 0.140, *t*(443) = 10.20, *p* < .001. Both relationships were nonlinear (weight scale: β = 0.273, *t*(443) = 11.58, *p* < .001; context strength: β = 0.101, *t*(443) = 6.45, *p* < .001). Critically, weight scale and context strength interacted, β = 0.193, *t*(443) = 11.33, *p* < .001, such that large transition costs were concentrated where both parameters were high.

Together, these sweeps revealed that lumping was the predominant pattern across the sampled parameter space. Most parameter combinations produced little or no transition cost, consistent with broad generalization across contexts. Substantial transition costs emerged only in a restricted region where high initial weight scale was paired with strong contextual signals. We therefore next asked whether these behavioral differences reflected corresponding changes in the networks’ internal representations.

### SRNs show that representational splitting requires both network flexibility and a strong context signal

To isolate the contributions of network scale and context strength, we selected representative low and high values from the parameter sweeps to form a 2 × 2 design (Figure 2C; see Methods for more details). This produced four conditions: Low-Scale + Low Context, Low-Scale + High Context, High-Scale + Low Context, and High-Scale + High Context. Networks were evaluated after 90 training sequences, matching the number of non-warmup sequences in the human experiment. If either factor alone were sufficient for splitting, High-Scale + Low Context or Low-Scale + High Context networks should exhibit transition costs (Figure 1D). If both were required, substantial costs should emerge primarily in the High-Scale + High Context condition.

All networks improved with training, although overall loss differed substantially across conditions (Figure 3A). These differences were partly expected from initialization: larger random weights in High-Scale networks produced more confident predictions, increasing cross-entropy when incorrect. By contrast, Low-Scale networks began with near-uniform predictions and loss close to chance.

**Figure 3.**
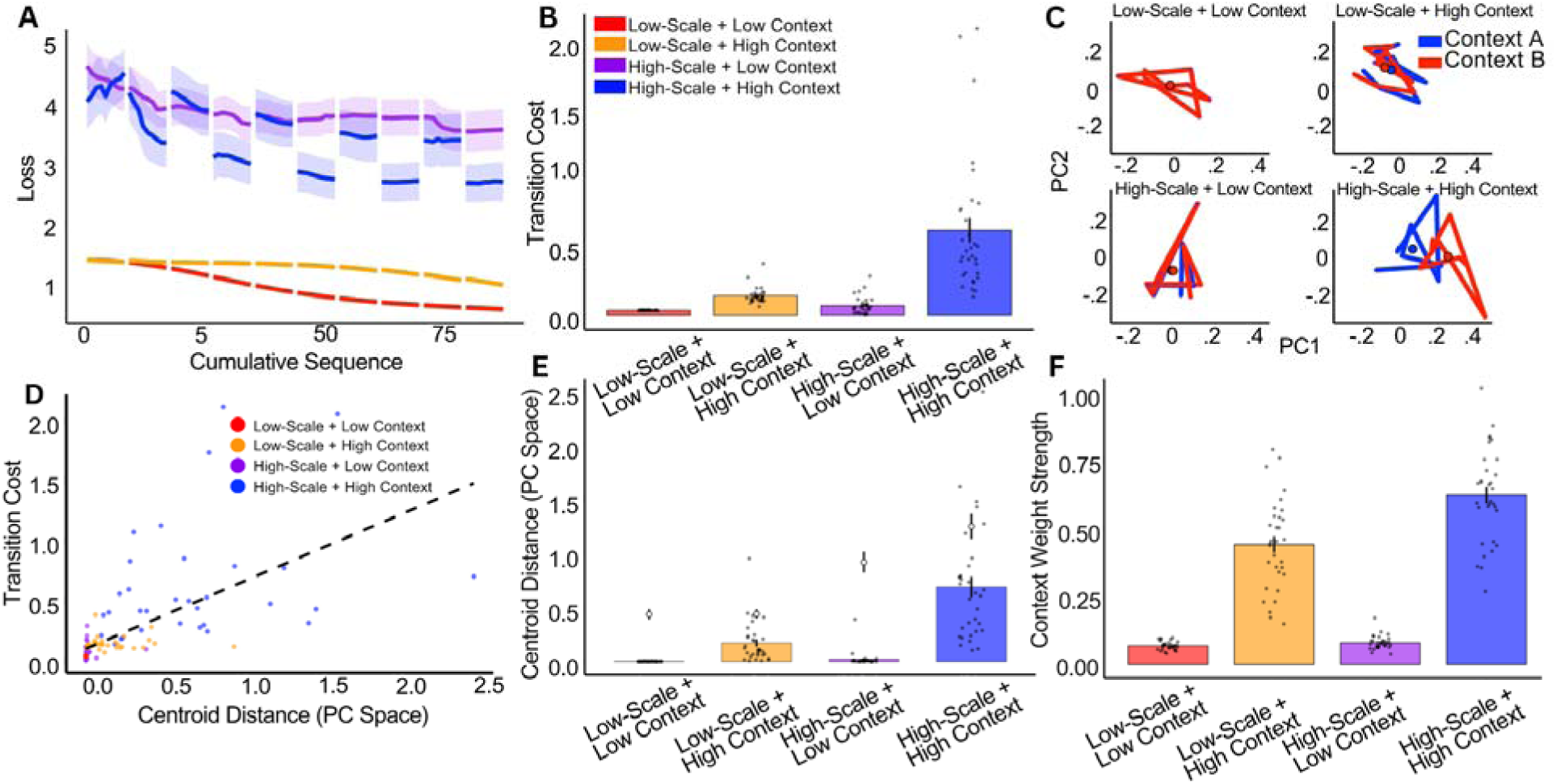
**(A)** Training loss across 90 sequences in the 2 × 2 design. Effects of network scale, *F*(1, 124) = 106.10, *p* < .001, η^2^ = .42, context strength, *F*(1, 124) = 16.45, *p* < .001, η^2^ = .06, and their interaction, *F*(1, 124) = 8.42, *p* = .004, η^2^ = .03. Shading indicates uncertainty across network runs. **(B)** Transition costs at the human-matched training endpoint were greatest with high network scale and strong context. Effects of network scale, *F*(1, 124) = 32.62, *p* < .001, η^2^ = .14, and context strength, *F*(1, 124) = 53.47, *p* < .001, η^2^ = .23, as well as their interaction, *F*(1, 124) = 24.26, *p* < .001, η^2^ = .10. **(C)** Mean hidden-state trajectories for Contexts A and B projected onto the first two principal components. Separation was greatest in the High-Scale + High Context condition. **(D)** Across network runs, greater Context A-B centroid separation was associated with larger transition costs. **(E)** Centroid distances within dual-context networks (colored bars) versus independently trained single-context networks (open circles), aligned b Procrustes analysis. Within-network distances remained below the independent-network benchmark. **(F)** Relative context-to-sequence input weight strength. Effects of network scale, *F*(1, 124) = 20.44, *p* < .001, and context-signal scale, *F*(1, 124) = 460.13, *p* < .001, as well as a network scale × context-signal scale interaction, *F*(1, 124) = 16.05, *p* < .001. Points in **B**, **E**, and **F** show individual network runs; bars and vertical lines show condition means and confidence intervals.

Having established that all conditions learned the sequence, we next asked whether switching contexts disrupted performance. Transition costs were greatest in the High-Scale + High Context condition, with a significant interaction between network scale and context strength (Figure 3B). Thus, the behavioral signature of splitting emerged most strongly when a strong context signal was available to a High-Scale network.

### Context-dependent costs reflect separation in hidden-state geometry

We next asked whether this behavioral pattern reflected corresponding differences in the networks’ internal representations. We quantified splitting as the Euclidean distance between Context A and Context B trajectory centroids in principal-component space (Figure 3C, E). Context separation showed main effects of network scale, *F*(1, 124) = 28.56, *p* < .001, η^2^ = .19, and context strength, *F*(1, 124) = 69.77, *p* < .001, η^2^ = .36, qualified by a network scale × context signal strength interaction, *F*(1, 124) = 24.57, *p* < .001, η^2^ = .17. The High-Scale + strong context-signal networks showed substantially greater Context A–Context B separation than the other conditions.

We also examined how networks weighted contextual information (Figure 3F). Strong-context networks placed greater weight on contextual inputs, yet Low-Scale + High Context networks showed limited representational separation and transition costs. Thus, greater context weighting alone was insufficient for splitting.

We also examined how transition costs were affected by learning rate and network size (Supplementary Figure 1) and also observed that these patterns persisted at matched levels of loss (Supplementary Figure 2) and after extended training (Supplementary Figure 3).

Importantly, greater Context A–Context B centroid separation was associated with larger transition costs across networks, *r*(126) = .63, *p* < .001 (Figure 3D), linking representational splitting to behavioral disruption at context switches.

We next asked how far this separation went. Specifically, did the most strongly split networks approach the separation expected if Context A and Context B were learned as fully independent solutions? To establish this benchmark, we compared within-network Context A–Context B distances with distances between independently trained Context A-only and Context B-only networks after rigid Procrustes alignment (see Methods). Across the four SRN conditions, distances were larger between independently trained single-context solutions than between Context A and Context B within the same network, *F*(1, 248) = 126.00, *p* < .001, η^2^ = .34. This effect varied by model type, *F*(3, 248) = 7.72, *p* < .001, η^2^ = .09, indicating that the gap between within-network and single-context distances differed across conditions. Critically, even the High-Scale networks, which showed the greatest within-network separation, remained less separated than independently trained Context A-only and Context B-only networks. Thus, High-Scale + High Context SRNs did not learn two fully independent representations.

### Humans show the behavioral cost predicted by context-dependent splitting

Having established the behavioral and representational signatures of splitting in SRNs, we next asked whether humans show a corresponding context dependence when learning the same motor sequence. Participants practiced the same eight-item finger sequence across 100 sequences of training. In the Single-Context conditions, participants practiced exclusively within either No Context, Maze or Melody contexts. In the Two-Context condition, participants alternated between Maze and Melody across blocks (Figure 4A). Critically, the required keypress sequence was identical across all conditions and remained visible throughout training. Context was therefore superfluous to successful sequence execution.

**Figure 4.**
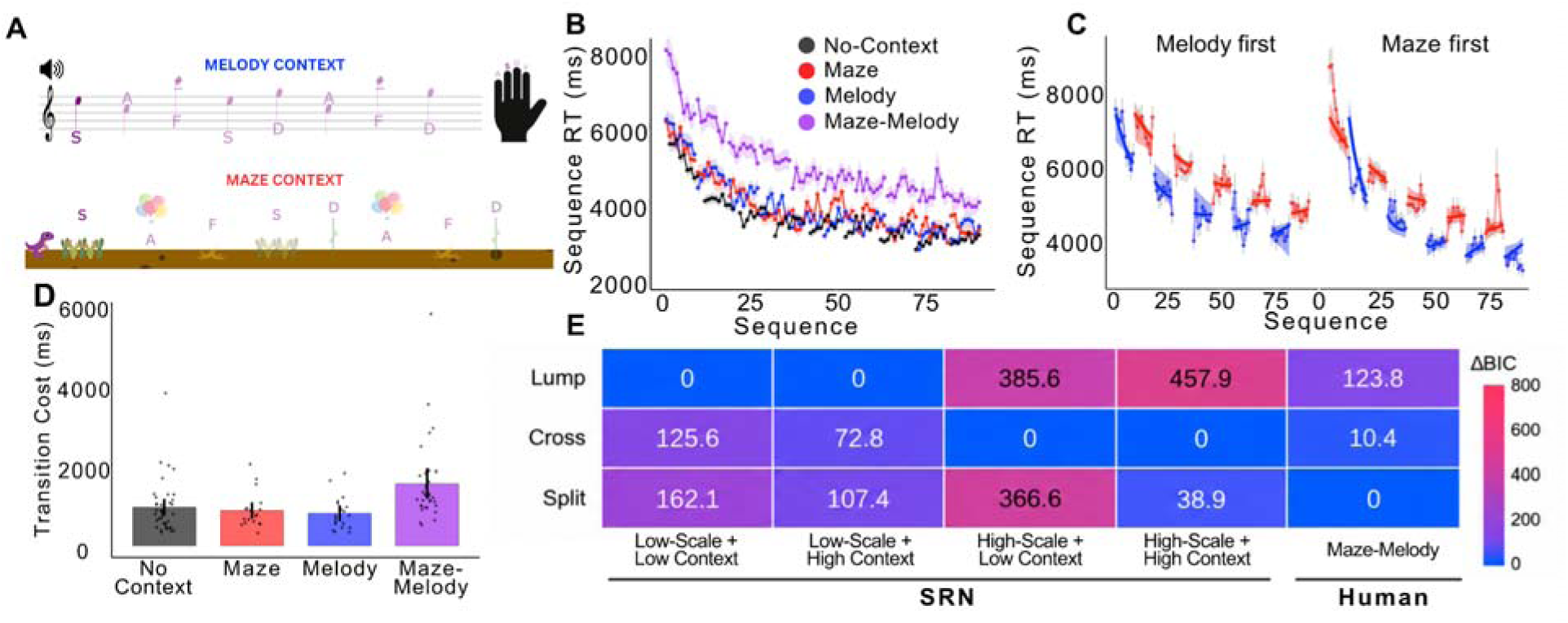
**(A)** Human task design. Participants practiced the same eight-item motor sequence in performance-inconsequential contexts. In the Melody context, sequence elements appeared as musical notes accompanied by tones; in the Maze context, responses advanced a character through a visual maze. Participants practiced without context, within a single Maze or Melody context, or while alternating between contexts. **(B)** Mean sequence response time across practice. All groups improved, although the Maze–Melody group remained slower than the single-context and no context groups. Shaded region indicate uncertainty around condition means. **(C)** Context-specific learning curves in the Two-Context group. Red and blue points show observed RTs in Maze and Melody, respectively. Smooth curves show the model fits from the best fitting split state-space model, capturing context-specific learning across alternating blocks. **(D)** Mean transition costs across block boundaries. Costs were larger in the Two-Context condition than in the single-context conditions. Points represent individual participants; bars and vertical lines show means and 95% confidence intervals. **(E)** State-space model comparisons for SRN and humans. Cells show ΔBIC relative to the best-fitting model within each condition, with lower values indicating better fit. Low-Scale SRNs favored the lumped model, whereas High-Scale SRNs favored the cross-talk model. Human Maze–Melody learning favored the split model.

The lumped and split accounts make different predictions for the shape of learning in the two-context condition. If learning fully generalized across contexts, practice in Maze and Melody should update a common representation, producing a relatively smooth learning trajectory across context changes. If learning became context dependent, however, performance should instead become organized by the current context, producing transition costs when participants moved between Maze and Melody. Across acquisition, sequence response time (RT) decreased reliably with practice for all groups, *F*(1, 9441.2) = 4163.77, *p* < .001, η*^2^ = .25* (Figure 4B). The two-context group was nevertheless slower overall than the other groups, *F*(1, 108) = 13.18, *p* < .001, η*_p_^2^ = .109*. Importantly, this difference was apparent early on in training, before repeated context alternation could produce an accumulated failure of transfer. We therefore do not interpret the overall performance difference itself as evidence for splitting. One possibility is that the two-context condition imposed additional task-set or contextual demands from the outset, but the present design cannot determine the source of this initial difference. The group difference changed over practice, reflected in a Sequence × Group interaction, *F*(1, 9441.2) = 297.86, *p* < .001, η*_p_^2^ = .02*. In contrast, performance did not differ reliably between the Single-Context groups, *F*(2, 78.5) = 0.85, *p* = .431, η*^2^ = .021*, indicating that simply embedding the sequence within either context did not impose a general performance cost. Training error rates also did not differ reliably across groups (Supplementary Figure 5), suggesting that the RT differences were unlikely to reflect a speed–accuracy trade-off.

More diagnostic was the structure of learning within the two-context condition. When Maze and Melody trials were examined separately, performance did not follow a single smooth trajectory across training. Instead, alternating contexts produced distinct, context-linked segments of the learning curve in both counterbalance orders (Figure 4C). Within the two-context condition, Melody was slower than Maze in both counterbalances, although the magnitude of this difference depended on which context participants experienced first (Supplementary Figure 6). Thus, even though participants repeatedly executed the same motor sequence, performance depended on which context currently framed that sequence. This pattern is the behavioral signature predicted by a split rather than fully lumped representation.

We next quantified these discontinuities at the boundaries between blocks, like we did for the SRNs. The Single-Context conditions provided a baseline for the change expected across an equivalent break without a change in context, whereas boundaries in the two-context condition additionally involved switching between Maze and Melody. Transition costs differed across conditions, *F*(3, 104) = 6.02, *p* < .001, η^2^ = .15 (Figure 4D). Critically, the two-context group showed substantially larger transition costs than the pooled single-context Maze and Melody groups (1,541 vs. 841 ms), *t*(40) = 3.61, *p* < .001, *d* = 0.89. Performance therefore changed more sharply across a block boundary when the context also changed than when participants resumed the same context after an equivalent break.

### State space modeling suggests both humans and high-scale SRNs split representations

To ask whether the context-dependent learning observed in humans and SRNs reflected similar underlying learning dynamics, we fit both systems within a common state-space framework (Verstynen et al., 2012; Gupta & Rickard, 2024). The same linear decoder mapped the currently expressed learning state onto the observed measure in both systems: human response times (RT) and SRN prediction loss. This approach allowed us to move beyond the discontinuities visible in the learning curves and directly test how much learning in one context transferred to the state governing the other under the same alternative learning-state architectures. Because human RTs also contain within-trial slowing and recovery across breaks (Rickard et al., 2008; Gupta & Rickard, 2022, 2024; Das et al., 2025), the human models additionally included a fatigue process to separate these transient performance changes from the slower learning dynamics of interest (see Methods). Stepwise model comparisons showed that learning was essential for capturing single-context performance, whereas the contributions of fatigue accumulation and recovery varied across conditions (Supplementary Table 1).

Let ***x_n_*** denote the latent learning state on sequence ***n***, and ***A*** the proportion of that state retained after each sequence. Learning was modeled using recursive state updates, with smaller values of ***A*** producing faster learning-related changes. The lumped model (M1) assumed that experience in either context updated a single latent learning state:

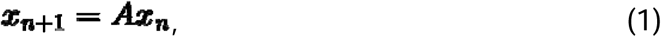

Because both contexts depended on the same state, learning acquired in one context necessarily carried over to the other. This model therefore represented complete lumping.

The cross-talk model (M2) instead contained separate latent states for Context A and Context B. Each state had its own update parameter, or , and could also be influenced by the state governing the other context. When Context A was active,

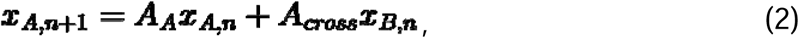

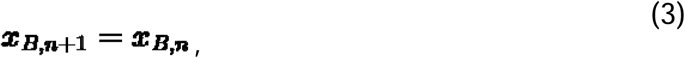

and when Context B was active,

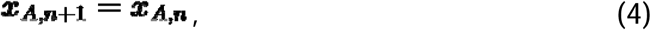

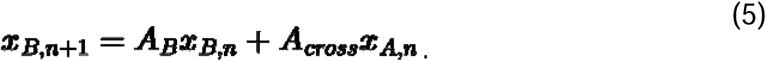

Thus, only the active context’s state was updated, while the inactive state remained unchanged. The parameters and controlled how much of each state’s existing contribution was retained during an update, while the cross-talk parameter ***A_Cross_*** controlled how strongly the other context’s state influenced its update. Larger values of ***A_Cross_*** indicated greater coupling between contexts, whereas values approaching zero indicated increasingly independent learning.

Finally, the fully split model (M3) was the limiting case of the two-state model in which ***A_Cross_ = 0***. When context A was active,

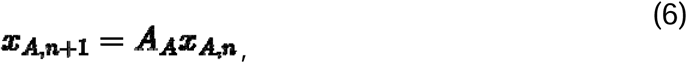

and when Context B was active,

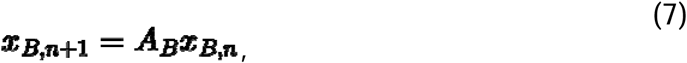

while the inactive state was held fixed as in equations 3 and 4. M3 therefore represents complete context-specificity: practice in Context A does not update the latent state governing Context B.

For both humans and SRNs, the context active on sequence ***n*** determined the latent learning state 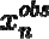 supplied to a common linear decoder. For SRNs,

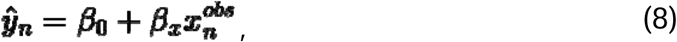

where ***y_n_*** was a prediction loss. For humans, we retained the same learning-state decoder but added the observable fatigue state 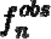 ,

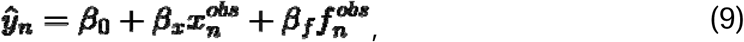

where ***y_n_*** was sequence RT. Based on prior work (Gupta & Rickard, 2022, 2024; Das et al., 2025), fatigue was assumed to accumulate within a trial and recover across breaks, allowing the model to separate transient performance slowing from the slower learning dynamics of interest. Thus, the comparison among lumped, coupled, and split learning states was the same for humans and SRNs; the human model contained an additional fatigue process needed to capture the structure of RT.

In the human single-context conditions, we fit only a single-state model, because models with separate context-specific states are not meaningful when only one context is present. We used the Single-Context conditions to determine which state-space components were necessary to capture ordinary learning dynamics, including learning, fatigue accumulation, and recovery (Supplementary Table 1).

We then carried this common nuisance structure into the Maze–Melody analysis and compared alternative learning-state architectures. When participants alternated between Maze and Melody, the fully split model provided the best fit (BIC = 2708), outperforming both the lumped model (BIC = 2832; ΔBIC = 124) and the cross-talk model (BIC = 2719; ΔBIC = 10.4). Thus, the architecture comparison was restricted to the multi-context condition, where separate latent states are interpretable, and favored distinct context-specific learning states over a shared state.

The SRNs showed a corresponding dependence on network scale and context strength. Low-Scale and High-Scale + Low Context networks were best captured by the lumped model (BICs = −1211 and −912, respectively), whereas Low-Scale + High Context networks were best captured by the cross-talk model (BIC = −1518; Figure 4E; complete model comparisons in Supplementary Table 2). High-Scale networks were also best captured by the cross-talk model (BIC = −858), which outperformed both the split model (BIC = −819; ΔBIC = 38.9) and the lumped model (BIC = −400; ΔBIC = 458).

However, the estimated cross-talk parameters were quite small in High-Scale networks (Maze-first: A_Cross_ = 0.031, 95% CI [0.000, 0.148]; Melody-first: A_Cross = 0.016, 95% CI [0.000, 0.098]). In the Low-Scale + High Context condition, cross-talk was slightly larger and more stable (Maze-first: A_Cross_ = 0.063, 95% CI [0.058, 0.071]; Melody-first: A_Cross_ = 0.054, 95% CI [0.048, 0.062]), but remained small. Thus, the SRN conditions showing the strongest context dependence were better described by weakly coupled context-specific states than by either a single shared state or fully independent states. All fitted parameters and recovery results for humans and SRNs are reported in Supplementary Tables 3 and 4, respectively.

Together, these results suggest that context strength and network scale jointly determine the degree of state separation. Low Context networks were adequately captured by a shared state, whereas stronger context dependence favored distinct but interacting latent states. In High-Scale networks, the strong BIC advantage of the cross-talk model over the lumped model, combined with relatively weak estimated coupling, suggests substantial context-specific separation without fully independent dynamics.

## Discussion

We asked whether the same motor sequence, practiced in different contexts, is learned through a shared representation or split into context-specific representations. Returning to the melody example from the Introduction, would practicing one song on the piano transfer to another song requiring the same finger movements, or would each song become a separate skill? Our SRN simulations revealed that lumping was the predominant solution across the parameter space, while splitting emerged when strong contextual signals were paired with high initial weight scale. Neither factor alone was sufficient. White-box analyses further showed that networks with greater separation between context-specific representations exhibited larger transition costs, despite learning identical sequences.

Perhaps more strikingly, humans showed a corresponding pattern. Participants practiced the same visible finger sequence in two contexts that provided no information necessary for successful performance. Nevertheless, alternating between Maze and Melody produced context-specific learning curves and larger transition costs than practicing within a single context. State-space modeling further favored split learning states in humans and weakly coupled states in the most context-dependent SRNs. Together, these findings show that shared task structure does not guarantee shared learning. The same sequence can become one skill or multiple context-specific skills, depending on the strength of contextual information and how the learning system organizes experience.

One way to understand this is to consider how learners weigh the features shared across experiences against those that differ. Every learning experience contains features from both the task being performed and the context in which it occurs. Two experiences may share identical task features while differing in contextual features, which can influence memory even when they are irrelevant to successful performance (Tulving & Thomson, 1973; Reder et al., 1974; Godden & Baddeley, 1975). Whether learning generalizes across experiences may depend on their subjective similarity, determined by the relative importance of shared and differing features (Tversky, 1977; Shepard, 1987; Tenenbaum & Griffiths, 2001; Marjieh et al., 2024). Models of categorization and context-sensitive semantic representations show how attention and inferred context can change the weighting of features, thereby altering psychological similarity (Nosofsky, 1986; Kruschke, 1992; Giallanza et al., 2026). However, similarity alone does not determine whether experiences update a shared representation or separate ones. Latent-cause inference provides a normative account: learners use observed features and their learned statistical relationships to infer whether experiences arise from the same underlying cause or different causes (Collins & Frank, 2016; Gershman et al., 2010; Gershman & Niv, 2010). Experiences attributed to a common cause can be learned together, whereas those attributed to different causes may be learned separately. Thus, even when task features are identical, the differences in contextual features may lead a learner to lump experiences into a shared representation or split them into context-specific representations.

This inference problem has received little attention in the domain of motor sequence learning. Here, learning is often characterized as a progression from stimulus-guided responding toward fluent, internally represented action sequences (Abrahamse et al., 2013; Doyon et al., 2009; Hikosaka et al., 2002). However, becoming less dependent on external stimuli does not necessarily mean that learning becomes independent of the context in which those actions are performed. Indeed, changing contextual cues can disrupt sequence performance, although these effects diminish with extensive practice (Ruitenberg, De Kleine, et al., 2012), are weaker for simpler sequences (Wright & Shea, 1991), and depend on which perceptual features are changed (Abrahamse & Verwey, 2008). These differences may reflect the degree to which contextual features are relevant or salient during learning and generalization across experiences. Supporting this possibility, recent work in motor adaptation found that emphasizing different contextual dimensions through task instructions changed how learned movements generalized across target shapes and directions (Poh et al., 2026). The contextual inference model offers a related computational account in which inferred contexts govern the creation, updating, and expression of motor memories (Heald et al., 2021). Together, these findings suggest that the relevance assigned to contextual features can shape generalization and the organization of motor learning. However, our SRN results indicate that contextual relevance alone may be insufficient to produce representational splitting. Initial weight scale had the strongest influence on transition costs across parameter sweeps. Learning rate also had an influence on transition costs. Network size, on the other hand, did little to induce splitting in Low-Scale networks (Supplementary Figure 1).

Beyond the importance assigned to contextual features, representational splitting also appears to depend on the organization of the learning system. In our SRNs, strong contextual inputs produced relatively overlapping representations at low initial weight scale but substantial separation at High-Scale. This is consistent with work demonstrating that initial weight scale influences how networks develop internal representations (Chizat et al., 2019; Dominé et al., 2025; Flesch et al., 2022; Holton et al., 2026; Woodworth et al., 2020). Although, the relationship between scale and splitting appears to be complex and likely depends on the task demands (Flesch et al., 2022; Holton et al., 2026).

Initial weight scale clearly plays a role in determining how learning is organized in the model, however, it remains unclear which properties of biological learning systems might serve an analogous role. One possibility is that it reflects the effective coupling of neural circuits, which is determined by both synaptic strength and connectivity structure. Supporting this possibility, neural network connectivity structure and its alignment with task statistics have been shown to influence learned representations independently of initial weight scale (Liu et al., 2024), while network topology alters how connection strength influences recurrent dynamics and learning (Mastrovito et al., 2024). Such differences in circuit organization may help explain why learners vary in their tendency to lump or split representations. Indeed, Holton et al. (2026) observed individual differences in human transfer and interference that resembled the behavior of networks with shared or distinct representations.

The organization of these circuits may reflect both inherited constraints and prior experience. Twin studies have identified genetic contributions to motor skill acquisition (Fox et al., 1996) and individual differences in brain network topology (Pourmotabbed et al., 2024). At the same time, the influence of experience on generalization has long been emphasized in stimulus-sampling and motor schema theories (Estes & Burke, 1953; R. A. Schmidt, 1975). Consistent with these accounts, the breadth of motor practice and variability in incidental context can influence subsequent generalization (Berniker et al., 2014; Apolinário-Souza et al., 2021). More recent work has demonstrated that prior training can reshape recurrent network dynamics and subsequent learning, as well as observing corresponding effects in biological neural circuits (Hocker et al., 2025; Bowler et al., 2026). Together, these findings suggest that the tendency to lump or split may depend on both inherited properties of neural circuits and their reorganization through experience.

Our SRN simulations identified that strong contextual signals and high initial weight scale were jointly necessary for splitting. However, our human experiment could only test the role of context, as the biological counterpart of weight scale remains unclear. Identifying such a counterpart may require examining task characteristics, individual differences, and prior training. Manipulating context also presents a challenge because contextual changes can affect performance independently of how learning is represented. Participants in the two-context condition were slower from the beginning of training (Figure 4A), potentially because they anticipated alternating between Maze and Melody. This resembles the mixing cost observed when participants perform multiple tasks within a block rather than a single task (Monsell, 2003; Rubin & Meiran, 2005). Mixing costs have been linked to sustained cognitive control demands and competition between task sets (Braver et al., 2003; Rubin & Meiran, 2005). Although Maze and Melody required identical motor responses, maintaining context-specific representations may have imposed similar demands. Transient switching costs offer another possible explanation (Rogers & Monsell, 1995). Our analyses excluded the first sequence of each trial (Gupta & Rickard, 2022; Gupta & Rickard, 2024; Supplementary Figure 4), largely eliminating the influence of immediate switching costs. A more comprehensive account must therefore separate the cognitive demands of anticipating and experiencing contextual switches from changes in the underlying representations.

Nevertheless, our findings suggest that the boundaries of a learned skill are not determined by its motor demands alone (Gandolfo et al., 1996; Howard et al., 2013; Wiestler et al., 2014) but also by higher-level context (Abrahamse & Verwey, 2008; Poh et al., 2016; Yokoi & Diedrichsen, 2019) and how the brain organizes experience (Holton et al., 2026). Returning to the budding pianist, practicing a sequence of notes in one song may not transfer seamlessly to the same sequence in another, even when the required movements are identical. Whether that practice generalizes depends on whether the surrounding contexts become part of what is learned. Thus, the same movements need not constitute the same skill.

## Methods

### SRNs

The goal of this study was to determine when learning the same sequence under different contexts gives rise to a shared representation (lumping) or context-specific representations (splitting). We leveraged SRNs to gain access to the internal representational structure, which cannot be as easily inferred from behavior alone. Critically, we focused on learning of the same sequence, which at first glance should have a lumped representation and consequently learning should generalize across contexts, thus providing the most stringent test of network architecture and context. Each network consisted of a single recurrent layer with six tanh units followed by a linear output layer over a four-item vocabulary. At each time step, the network received the current sequence item together with a two-dimensional one-hot context vector and predicted the next item in the sequence. The recurrent state therefore evolved according to the current sequence input, the context signal, and the preceding hidden states.

Sequence items were represented using Gaussian-smoothed vectors rather than discrete one-hot vectors. This introduced graded overlap between neighboring finger representations, approximating the distributed and correlated neural activity patterns associated with individual finger movements in sensorimotor cortex (Ejaz et al., 2015).

For a token □, activation across the four input dimensions decreased as a Gaussian function of distance from □, with bandwidth □ = *0.7*, and the resulting vector was normalized to sum to one. This encoding introduced partial similarity among neighboring sequence items while preserving the identity of each item. The context vector was concatenated with the token representation at every time step and supplied directly to the recurrent layer. Networks were trained by minimizing cross-entropy loss using the Adam optimizer with a learning rate of .005.

### Parameter sweeps

Before focusing on the four representative conditions that crossed initial weight scale and context-signal scale, we mapped transition costs across a broader parameter space. The weight-scale sweep varied weight initialization scale while holding context-input scale stable; the context strength sweep varied context-input scale at both Low-and High-Scale network settings, and the joint sweep crossed both parameters. The one-dimensional sweeps sampled 30 logarithmically spaced values for weight scale (.01–4.0) and context strength (.005–8.0). The joint sweep sampled a 15 × 15 logarithmic grid. The four parameter combinations used in the primary 2 × 2 design were embedded within this broader parameter landscape and were selected to be representative samples from that landscape. Networks in the parameter sweeps were trained for 500 sequence repetitions arranged in alternating 10-sequence Context A and Context B blocks.

### Network scale and context strength

We manipulated two properties of the SRNs at initialization: network scale and context strength. Network scale controlled the initial magnitude of the sequence-input, recurrent, and output weights, which were sampled from zero-mean Gaussian distributions with standard deviations of 0.02 (Low-Scale) or 2.0 (High-Scale). Context strength independently controlled the magnitude of the input-to-hidden weights connecting the context vector to the hidden layer, with standard deviations of 0.02 (Low Context) or 2.0 (High Context). All biases were initialized to zero.

These manipulations allowed us to vary the network’s initial dynamics and the influence of contextual information separately. Network scale altered the initial magnitude of the sequence-input and recurrent weights, whereas context strength controlled the initial strength of the connections carrying contextual information into the hidden state. We refer to the latter’s influence as context signal strength. Because the networks used tanh activation functions, larger initial weights could also produce more nonlinear or saturated responses (Glorot & Bengio, 2010; Sutskever et al., 2013). Crossing these parameters produced four conditions: Low-Scale + Low Context, Low-Scale + High Context, High-Scale + Low Context, and High-Scale + High Context. Importantly, context strength changed the weights connecting context to the hidden layer, not the numerical values of the context vectors themselves.

Each network was trained on the same fixed eight-item sequence in Context A and Context B. Because the input sequence and next-item targets were identical across contexts, context provided no information required to solve the prediction problem. The network could therefore achieve low prediction loss while either representing the two contexts similarly or developing distinct context-specific hidden states.

For the human-matched analyses, networks followed the same sequence of training contexts and observations as the corresponding human participant. Each participant × model-condition pair was initialized independently, while the same random seed was used across the four model conditions for a given participant so that differences among conditions reflected the parameter manipulations rather than different random initializations. Networks were evaluated at the human-matched training endpoint for the primary analyses and subsequently trained further to examine whether representational differences persisted with extended training.

### SRN white-box analyses

Access to the SRNs’ hidden states and weights allowed us to examine how context-dependent representations related to behavioral transition costs. We focused on two complementary measures: separation between context-specific hidden-state trajectories and the relative strength of the learned context-input weights.

### Hidden-state trajectories and context separation

For each trained network, we presented the same eight-item sequence in Contexts A and B and recorded the six-dimensional recurrent hidden state at each sequence position. We projected the hidden states from both contexts onto the first two principal components using a shared PCA transformation. These trajectories provided a visualization of how the network represented the same sequence across contexts, with greater separation indicating more differentiated hidden-state activity.

To quantify this separation, we calculated the centroid of each trajectory by averaging its PC1 and PC2 coordinates across sequence positions. Context separation was defined as the Euclidean distance between the two centroids:

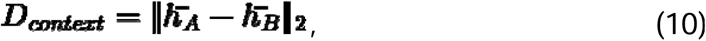

where 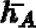 and 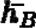 are the two-dimensional trajectory centroids for Contexts A and B. Smaller values indicate that the two contexts occupy similar regions of the projected hidden-state space, whereas larger values indicate greater separation.

We also compared this within-network separation against a benchmark derived from independently trained Context A-only and Context B-only networks. Because independently trained networks can have differently oriented representational spaces,

we used Procrustes alignment to place their trajectories in a common coordinate system (Goodall, 1991). For each model condition, we first averaged the projected trajectories across independently trained networks at corresponding sequence positions. We then estimated a Procrustes rotation and translation that aligned the mean Context B-only trajectory to the mean Context A-only trajectory and applied this transformation to the Context B-only networks. Finally, we calculated the centroids of individual single-context networks and compared distances between independently trained Context A-only and Context B-only networks with the within-network distances from the two-context condition (Figure 3E). This provided a reference for evaluating how closely context-dependent representations approached the separation observed between independently learned solutions.

### Context-weight strength

We also examined whether representational separation was accompanied by greater reliance on contextual information. We partitioned the input-to-hidden weight matrix into sequence-token weights, , and context-input weights, , and calculated their Frobenius norms, defined as the square root of the sum of squared matrix entries (Golub & Loan, 2013). Relative context-weight strength was defined as

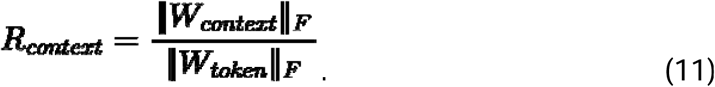

Larger values indicate stronger context-input weights relative to sequence-token weights. This measure allowed us to distinguish how strongly networks weighted contextual information from how extensively that information separated their hidden-state representations.

### Participants

One hundred and eleven (age = 27.41, F□=□46.3%) participants were recruited from the online Prolific participant pool in exchange for payment. All participants were 18 years of age or older, reported to be right-hand dominant, and provided informed consent via button press. All procedures were approved by Princeton’s institutional review board.

### Human Task

Participants learned to repeatedly type an eight-item sequence using the ‘A’, ‘S’, ‘D’, and ‘F’ keys on their computer keyboard with their non-dominant left hand. The eight-item sequence was selected using an exhaustive generate-and-filter procedure to balance key frequency and transition statistics while excluding obvious regularities.

Each block consisted of 10 sequences followed by a 10 second break. The sequence was explicitly displayed on the screen throughout the block. At the start of the experiment, participants completed a short familiarization in which they were required to execute two consecutive sequences with at least 90% accuracy in under 4s. Correct responses were signaled by a visual change on the screen relevant to the context (Figure 4A). The main experiment then consisted of 10 blocks where each block had 10 sequences. Participants were randomly assigned to one of the four groups. For the no Context group, participants were tasked with simply pressing the computer key of the presented letter (Figure 4A). For the single context groups, participants were assigned to either a “maze” context or a “melody” context. In the maze context, the sequence unfolds as a 2D side-scrolling platformer game where each key press allows an avatar to successfully navigate obstacles (Maze group; Figure 4A). In contrast, in the melody context, the sequence appears as staff-like notation that scrolls across the screen and each key press is accompanied by an auditory note (Melody group; Figure 4A). A correct button press was indicated by the avatar moving to the next position or the note on the staff changing color. Finally, for the Two-Context group, the participants alternated between the maze and melody contexts. Critically, all groups performed the same sequence each block with the only difference being the context in which it was executed.

### Transition Cost

To quantify the cost associated with switching contexts, we measured the discontinuity in performance across trial boundaries. For SRNs, transition cost at boundary □ was defined as the absolute difference between loss on the first sequence of the new block and the final sequence of the preceding block,

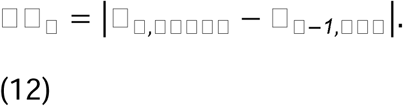

Transition costs were averaged across context switches within each network run to obtain a single estimate per SRN.

The same boundary-based measure was used for the human behavioral experiment, with sequence RT replacing loss. Because the first sequence of each human trial served as a warm-up and was excluded from analysis (Supplementary Figure 4), human transition cost was defined as the absolute difference between RT on Sequence 2 of the new trial and Sequence 10 of the preceding trial,

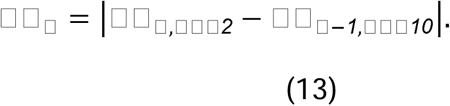

Transition costs were averaged across trial boundaries within each participant. In the Maze–Melody condition, these boundaries involved a context switch, whereas the single-context Maze and Melody conditions provided the corresponding baseline for trial-to-trial changes without a context switch.

### Human Behavioral Experiment Analysis

Participant-level exclusions were applied to remove low-quality data, which is common practice for studies conducted using online platforms (Crump et al., 2013; Masís Obando et al., 2025). First, we computed error rates and excluded participants whose error rate exceeded the group mean + 2.5 SD. Second, we excluded unusually slow responders using a two-pass criterion on participant mean RTs computed from correct, non-warmup trials: participants above mean + 2.5 SD were removed, the distribution was recomputed, and the same cutoff was applied again to define the final inclusion set. Sequence RT was defined as the sum of keypress RTs within each correctly completed sequence. Incomplete sequences due to an error keypress were removed. In humans, we removed the first sequence due to a large detriment to performance at the start of the trial, which eliminates the possibility that the transition costs are canonical switch costs for humans (Supplementary Figure 4; Gupta & Rickard, 2024, 2026).

### State-Space Model

We fitted state-space models to human RTs and SRN prediction loss to examine whether learning dynamics were better explained by shared or context-specific latent states. The three learning architectures (M1: Lump, M2: Cross-talk, M3: Split) and their transition equations are described in the Results. Here, we describe the preparation of the learning curves, the additional fatigue process included in the human models, and the procedures used for parameter estimation and model comparison.

### Fatigue

Human RTs reflect both learning-related improvements and transient slowing associated with reactive inhibition or fatigue (Rickard et al., 2008; Gupta & Rickard, 2022, 2024; Das et al., 2025). We modeled these processes separately to prevent within-trial slowing and recovery from being attributed to changes in the learning state.

A single latent fatigue state, ***F_n_***, was included in all three human model architectures. Fatigue was initialized at zero and accumulated across successive sequences within a block. Let ***C*** denote the fatigue-buildup parameter and ***m_b_*** the number of analyzed sequences in block ***b***. For successive sequences within the same block, fatigue increased according to

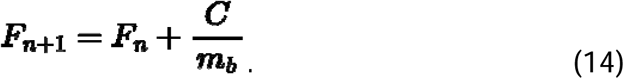

The increment was therefore constant within a block and scaled by the number of analyzed sequences. Because the increment was applied between successive observations, ***C*** represented a fatigue-buildup scale rather than the exact total increase across the observed block. Between blocks, fatigue recovered exponentially:

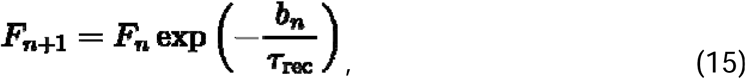

where ***b_n_*** was the break duration, set to 10 seconds for ordinary between-block breaks, and ***τ_rec_*** was the recovery time constant. Larger values corresponded to slower recovery, whereas values approaching zero corresponded to increasingly complete recovery during the break. The fatigue process was shared across contexts and model architectures. Consequently, differences in model fit reflected the organization of the learning states rather than differences in how fatigue was represented.

### Observation model and parameter estimation

For each candidate set of learning-state parameters, we simulated the latent trajectory and mapped the currently expressed state onto the observed RT or prediction loss using the linear observation models defined in the Results. Each fitted trajectory had one intercept and one learning-state coefficient, estimated by ordinary least squares. For the human models, the simulated fatigue state was added directly to the predicted RT with its coefficient fixed at one, placing fatigue in RT units and avoiding a scaling ambiguity between its buildup parameter and decoder coefficient.

The dynamic parameters were estimated by minimizing the residual sum of squares between observed and predicted values:

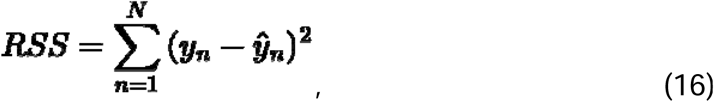

Where ***y_n_*** and ***ŷ_2_*** were the observed and predicted values for sequence ***n***. At each candidate parameter setting, we simulated the latent states, re-estimated the linear decoder coefficients, and calculated prediction error. The dynamic parameters were optimized using bounded L-BFGS-B optimization. Human fits used multiple initialization points to reduce sensitivity to local minima.

To preserve the nesting of the two-state architectures, the Cross-talk model was additionally evaluated from the best-fitting Split solution with its cross-talk parameter fixed initially at zero. This ensured that the more general Cross-talk model could achieve at least as good a fit as the Split model.

We also compared nested single-state models in the No Context condition to assess the contributions of learning, fatigue accumulation, and fatigue recovery. These comparisons included an intercept-only model, a learning-only model, a model with learning and fatigue accumulation under fixed recovery, and the full model with freely estimated recovery. The results are reported in Supplementary Table 1.

### Model fitting and comparison

The model was fit by minimizing the root mean squared error between observed and predicted RTs,

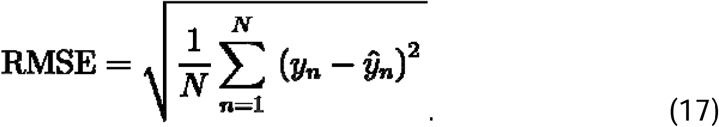

The dynamic parameters were optimized with bounded numerical optimization.

At each candidate parameter setting, the latent states were simulated and the decoder coefficients were re-estimated using least squares. This produced the best linear mapping from the simulated latent trajectory to the observed RTs for that parameter setting. We compared three model classes.

### Model comparison

Model fits were compared using BIC. We computed the residual variance as

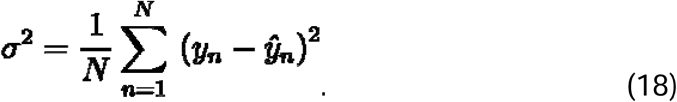

The Gaussian log likelihood was

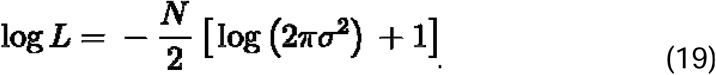

BIC was computed as

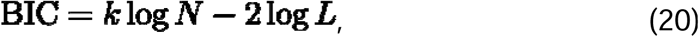

where *N* was the number of observations and *k* was the number of estimated parameters. For the human models, parameter counts included the learning-state parameters, fatigue parameters, two decoder coefficients, and residual variance. This yielded 6, 8, and 7 parameters for Lump, Cross-talk, and Split, respectively.

The alternating-context human models were fitted independently for each counterbalance. We then summed their log-likelihoods and parameter counts and calculated a combined BIC using the total number of observations across both counterbalances. Lower BIC values indicated a better balance between model fit and complexity.

For the SRN models, BIC was calculated from the residual-based expression ***N*log(*RSS/N*) + *k* log (*N*)**, using 3, 5, and 4 fitted parameters for Lump, Cross-talk, and Split, respectively. These counts included the dynamic parameters and two decoder coefficients. Models were compared within each dataset rather than by comparing absolute BIC values across humans and SRNs.

## Acknowledgements

We thank the members of the Intelligent Performance and Adaptation Laboratory (IPA Lab) in the Department of Psychology at Princeton University for many fruitful discussions. We thank Jonathan Cohen for his insights and broader perspective on neural networks and Eleanor Holton for her insights into how initial weight scale influences neural network function. This project was supported by the Innovation Fund for New Ideas in the Natural Sciences from the Office of the Dean of Research at Princeton University. MWG and JAT were also supported by grants R01NS131552 and R01NS134754 from the National Institute of Neurological Disorders and Stroke (NINDS) of the National Institutes of Health (NIH). The funders had no role in study design, data collection and analysis, decision to publish, or preparation of the findings.

