## Supplementary material for "Same Movements, Different Skills: How Context Shapes Learning in Humans and Neural Networks": supplemetary

### Effects of Learning Rate and Network Size on Transition Costs

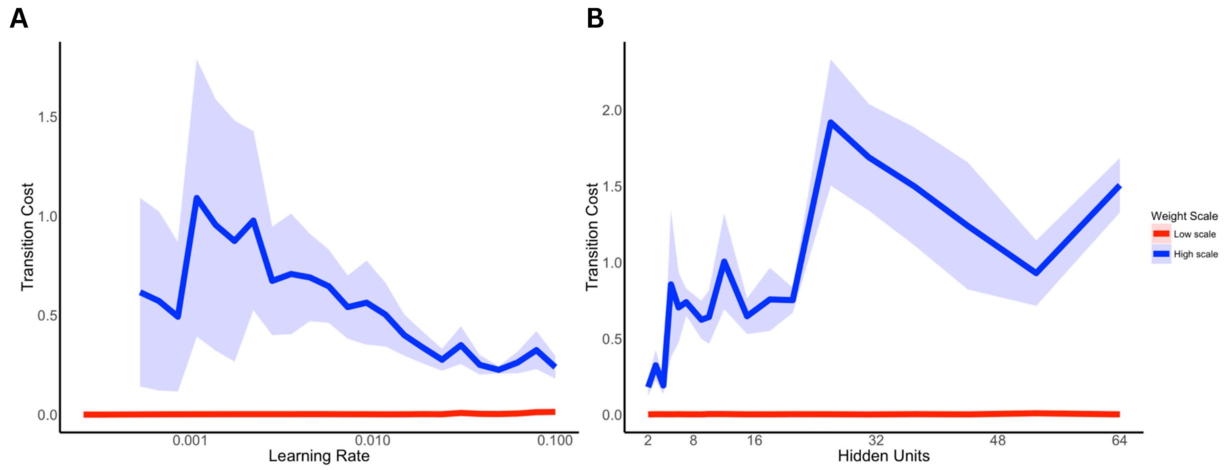


**Supplementary Figure 1. (**A) Transition costs across learning rates for Low-Scale + Low Context (blue) and High-Scale + High Context (red) networks. (B) Transition costs across network sizes, measured by the number of hidden units, for the same two network conditions. Lines indicate mean transition costs, and shaded regions represent ±1 standard error across network runs.

To examine how learning rate and network size influence context-dependent learning, we independently varied these parameters in Low-Scale and High-Scale SRNs (Supplementary Figure 1). We analyzed transition costs using linear mixed-effects models with fixed effects of network condition, the log-transformed parameter, and their interaction, with random intercepts for seed. All network runs were included. Adding quadratic terms did not improve model fit for either learning rate, $\chi^2(2) = 0.92$, $p = .632$, or network size, $\chi^2(2) = 0.23$, $p = .893$.

Both parameters influenced transition costs primarily in High-Scale networks, but in opposite directions. Increasing learning rate reduced transition costs in High-Scale networks, as indicated by a significant Learning Rate × Condition interaction, $\beta = -0.406$, $t = -7.23$, $p < .001$. In contrast, learning rate had no reliable effect in Low-Scale networks, $\beta = 0.003$, $t = 0.07$, $p = .946$. Increasing network size, however, produced larger transition costs in High-Scale networks, reflected in a significant Network Size × Condition interaction, $\beta = 0.263$, $t = 6.00$, $p < .001$. Again, the corresponding effect in Low-Scale networks was negligible, $\beta < .001$, $t = 0.01$, $p = .993$.

Together, these findings indicate that learning rate and network size modulate context-dependent transition costs, but neither produces substantial costs in Low-Scale networks under the tested conditions. Thus, while network architecture and learning dynamics influence the magnitude of context-dependent learning, their effects depend on the initial weight scale and contextual inputs.

**Matched Loss Analyses**

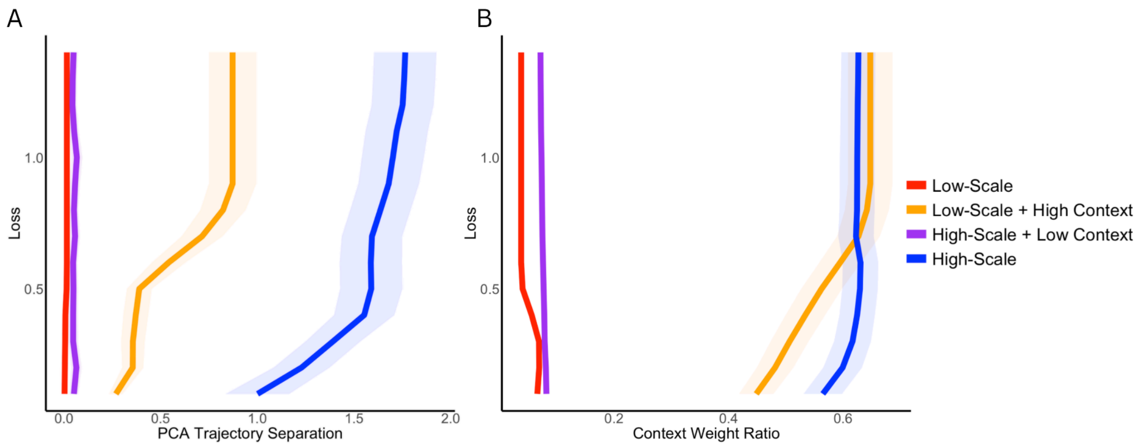


**Supplementary Figure 2.** Key representational differences persist at loss-matched checkpoints. To test whether differences between SRN conditions reflected greater learning progress rather than distinct representations, we compared networks at matched loss values. (A) PCA trajectory separation remained larger in the High-Scale condition than in the Low-Scale conditions across matched levels of loss. (B) Context/input weight ratios also differed across model conditions at matched loss values, with larger context weighting in the High-Scale and Low-Scale + High Context conditions. These patterns indicate that the mechanistic differences between conditions were not explained solely by on network learning differences.

To determine whether differences in context use simply reflected where networks were sampled along the loss trajectory, rather than the scale- and context-defined learning itself, we repeated the context-weight analysis on loss-matched checkpoints using a linear mixed-effects model with fixed effects of model type, centered matched loss, and their interaction, with random intercepts for network run. The Low-Scale + Low Context condition served as the reference group.

At the mean matched-loss value, the Low-Scale + Low Context networks assigned little weight to the context input, $\beta = 0.045$, $t = 3.94$, $p < .001$. Relative to this baseline, both high-context conditions showed substantially larger context-weight ratios: Low-Scale + High Context, $\beta = 0.547$, $t = 68.74$, $p < .001$, and High-Scale + High Context, $\beta = 0.575$, $t = 71.21$, $p < .001$. High-Scale + Low Context networks also showed a reliable but much smaller increase relative to baseline, $\beta = 0.030$, $t = 3.75$, $p < .001$. Thus, after matching networks on loss, context weighting was not explained by optimization progress alone. Instead, the strongest determinant of context use was the strength of the context signal itself.

The relation between matched loss and context weighting also differed across model types. In the Low-Scale + Low Context reference condition, context weighting showed only a marginal association with matched loss, $\beta = -0.025$, $t = -1.79$, $p = .074$. This slope was significantly more positive in the Low-Scale + High Context condition, $\beta = 0.181$, $t = 9.14$, $p < .001$, and modestly more positive in the High-Scale + High Context condition, $\beta = 0.043$, $t = 2.13$, $p = .033$. The High-Scale + Low Context condition did not differ reliably from the Low-Scale + Low Context baseline in its loss-dependent change in context weighting, $\beta = 0.017$, $t = 0.86$, $p = .388$.

These matched-loss analyses clarify the mechanism. Greater use of the context input was not merely a byproduct of some networks being further along in learning.

However, context weighting alone was also not sufficient to explain the full splitting effect: Low-Scale networks with a strong context signal assigned substantial weight to context, yet the broader behavioral and geometric signatures of splitting were strongest when a strong context signal was paired with high network scale. Thus, the updated results support a two-stage interpretation: strong context signals promote context use, but representational splitting emerges most clearly when that context signal is available to a sufficiently flexible network.

#### SRN Extended Learning

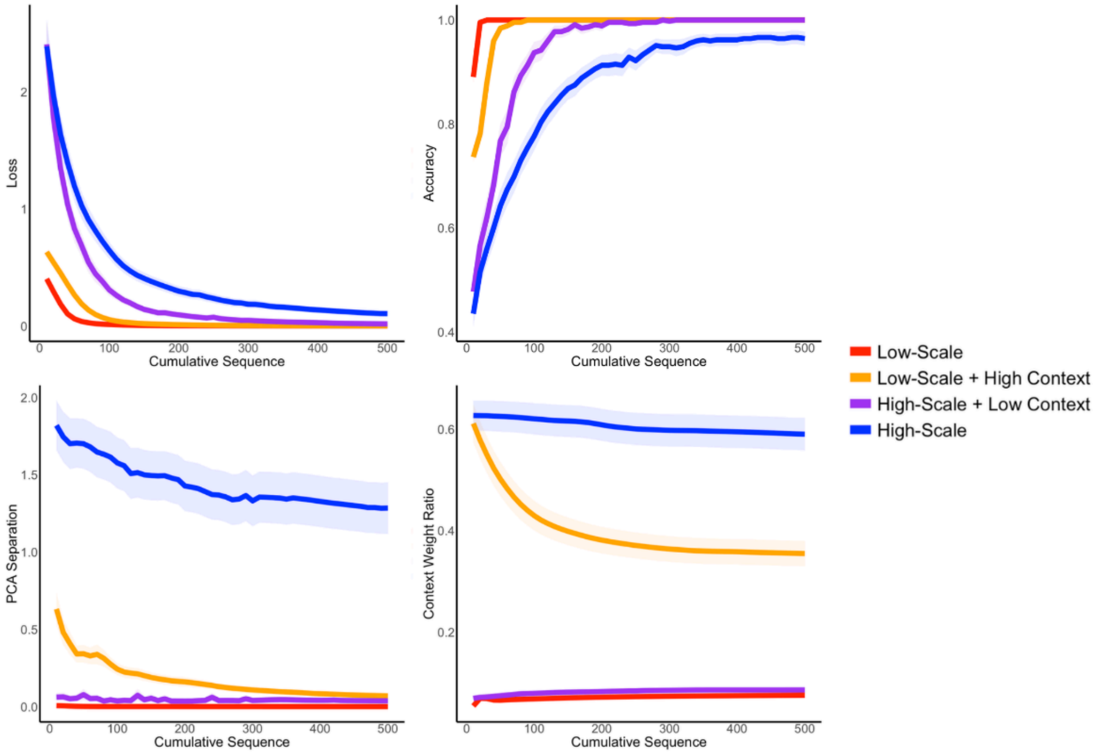


**Supplementary Figure 3.** Extended SRN training dynamics across model conditions. Loss, accuracy, PCA trajectory separation, and context weight ratio are plotted across 500 cumulative training sequences for each SRN condition. Lines show the mean trajectory for each model type, and shaded regions show uncertainty around the mean.

Generally, the patterns that were evident in the main text (Figure 3) hold across longer training.

To examine how each model metric changed across extended training, we fit linear mixed-effects models across 500 training steps. For each outcome, the model included fixed effects of model type, centered training step, and their interaction. We included random intercepts and random slopes for centered training step by network run, with a random-intercept-only fallback when the maximal random-effects structure was singular. The main effect of the training step tested whether each metric changed systematically over training. The main effect of model type tested whether the model conditions differed at the centered training step.

All metrics changed reliably across training. There was a main effect of training step for loss, $F(1, 128) = 286.84$, $p < .001$; accuracy, $F(1, 128) = 374.17$, $p < .001$; the context/input weight ratio, $F(1, 128) = 59.66$, $p < .001$; and trajectory separation, $F(1, 128) = 51.28$, $p < .001$. Thus, performance and representational metrics continued to evolve across the full 500-step training period.

Model type also affected most metrics. There were reliable main effects of model type for loss, $F(3, 128) = 47.39$, $p < .001$; accuracy, $F(3, 128.10) = 50.56$, $p < .001$; the context/input weight ratio, $F(3, 128.06) = 165.66$, $p < .001$; and trajectory separation, $F(3, 128.00) = 74.17$, $p < .001$. Thus, the model manipulations affected performance, context use, and trajectory separation across training.

Together, these analyses show that the SRN conditions differed across extended training in both task performance and internal representational structure. The strongest differences appeared in loss, accuracy, context/input weighting, and trajectory separation.

#### First sequence costs

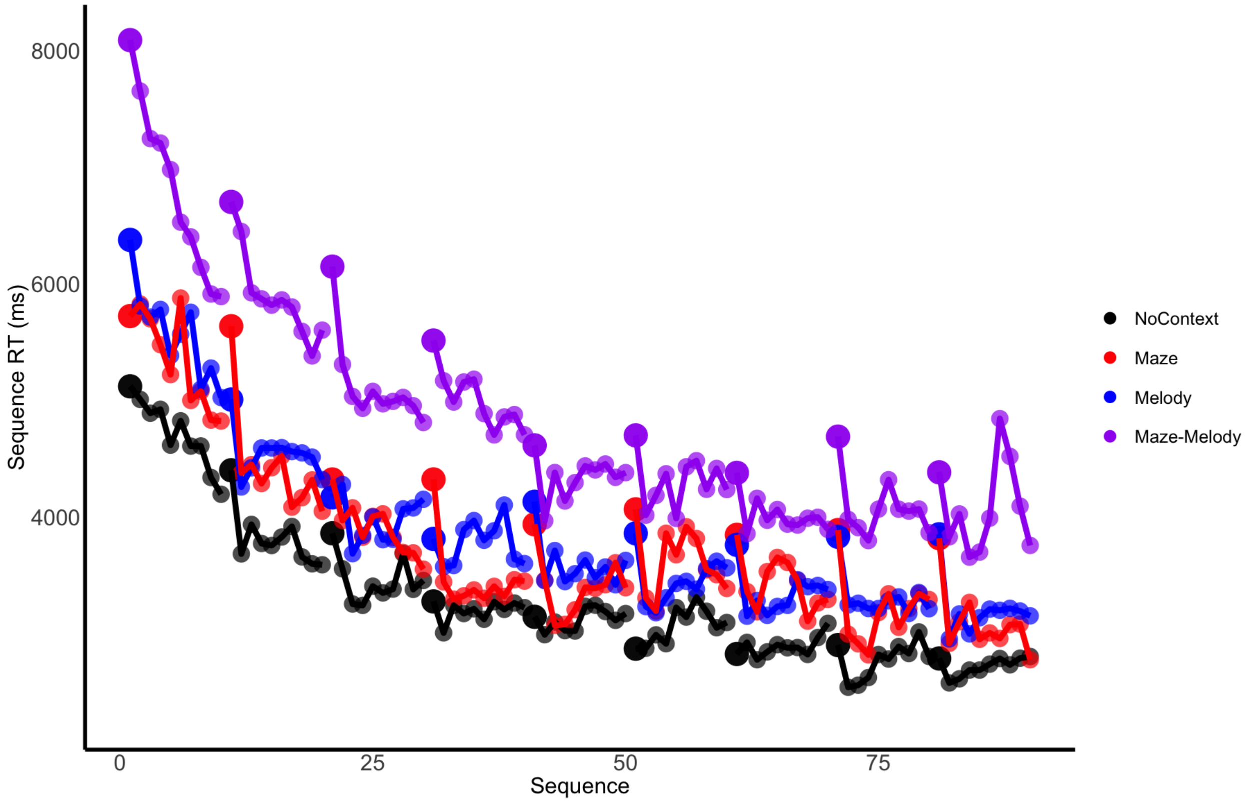


**Supplementary Figure 4.** First-sequence slowing across training. Each dot corresponds to mean sequence RTs across training for the No Context, Maze, Melody, and Maze-Melody groups. Larger points mark the first sequence of each trial. As visually evident in the graph, these sequences are generally much slower than other sequences within the trial for all groups, hence why they were removed from the analyses.

We quantified first-sequence costs for each participant as the difference between the first and second sequence within each trial (Gupta & Rickard, 2022; 2024). Specifically, we computed the cost as sequence 0 RT minus sequence 1 RT. Across groups, the first sequence was slower than the second sequence (sequence 0: 4071 ms; sequence 1: 3602 ms), producing an average first-sequence cost of 468 ms. A repeated-measures ANOVA confirmed a reliable first-sequence effect, $F(1, 107)$ =

66.12, $p < .001$, $\eta^2 = .02$. Thus, participants showed robust slowing at the start of each trial.

We next asked whether this cost was larger in the Maze-Melody group than in the single-context groups. The Maze-Melody group showed a numerically larger first-sequence cost than the combined comparison groups (Maze-Melody: 565 ms; single-context: 428 ms), but this difference was not reliable, $F(1, 106) = 1.18$, $p = .280$, $\eta^2 = .01$. The individual group means followed the same pattern descriptively: No-Context = 233 ms, Maze = 705 ms, Melody = 541 ms, and Maze-Melody = 565 ms. Thus, although first-sequence costs were present overall, the Maze-Melody group did not show a reliably larger first sequence cost.

**Error Analysis**

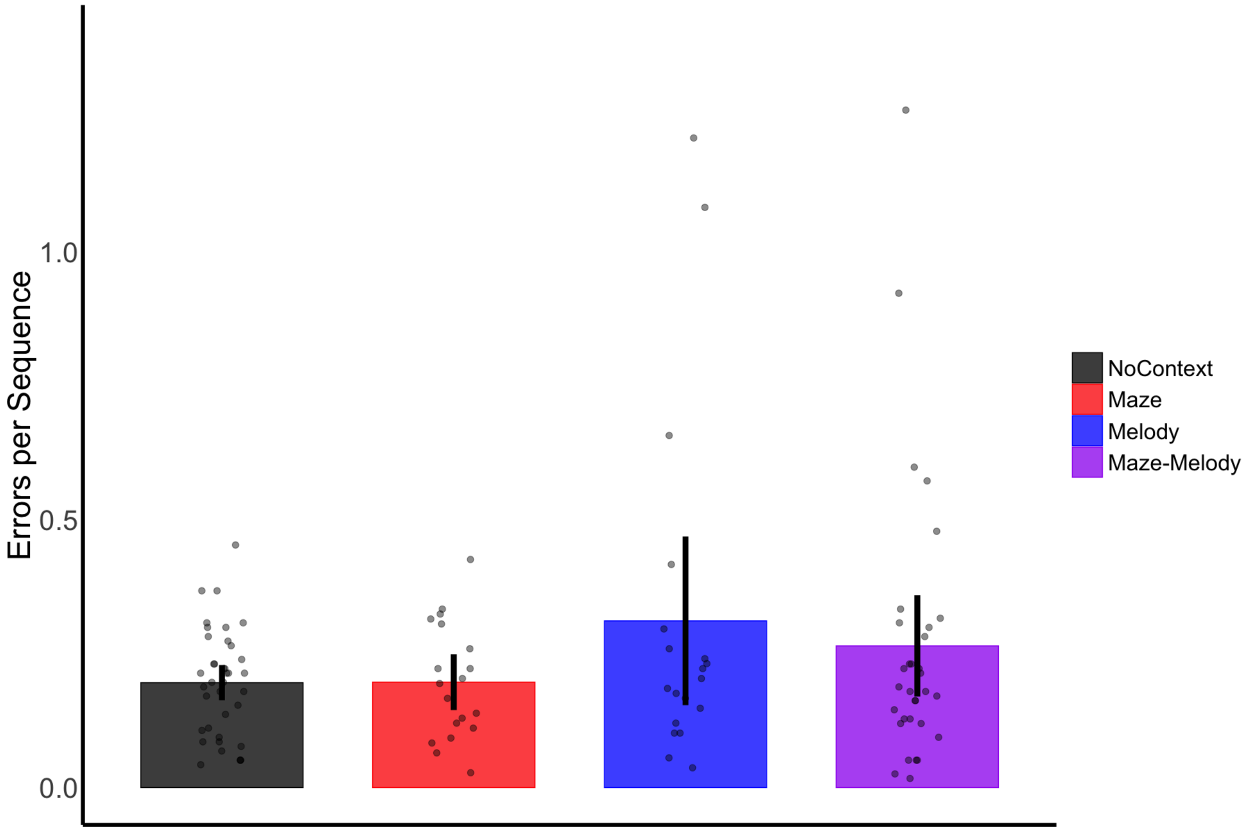

**Supplementary Figure 5.** Training errors by group. Bars show the mean number of incorrect keypresses made before completing a correct sequence within a trial. Points show participant-level means. Error rates were similar across the No-Context, Maze, Melody, and Maze-Melody groups, indicating that the training RT differences were unlikely to reflect a simple speed–accuracy trade-off.

We quantified errors for each participant as the number of incorrect keypresses made before completing a correct sequence within a trial. During training, the groups made similar numbers of errors (No-Context: 0.20; Maze: 0.20; Melody: 0.31; Maze-Melody: 0.27 incorrect keypresses per sequence). A one-way ANOVA showed no reliable difference between groups, $F(3, 104) = 1.68$, $p = .177$, $\eta^2 = .05$. Thus, the training RT differences are unlikely to reflect a simple speed–accuracy trade-off.

### Ordering effects

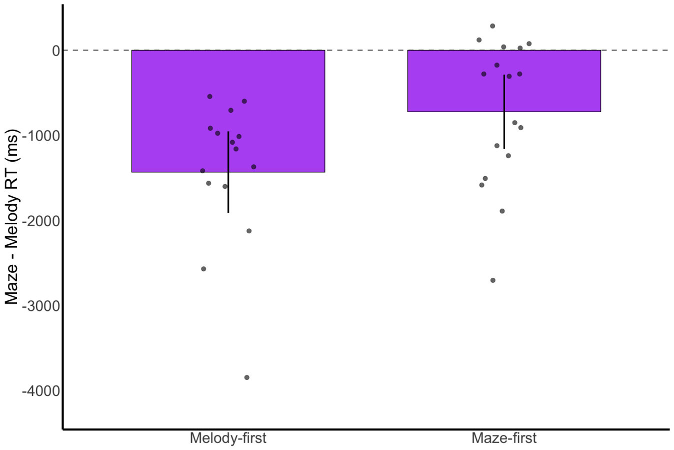


**Supplementary Figure 6.** (A) Start order modulates the size, but not the direction, of the context difference. Bars show the mean Maze minus Melody RT difference (in the Maze-Melody group) for participants who experienced Melody first or Maze first. Points show participant-level difference scores. Values below zero indicate that Melody trials were slower than Maze trials. Melody was slower than Maze in both start-order groups, but the Maze–Melody difference was larger for participants who experienced Melody first.

We next asked whether the order in which participants experienced the two contexts changed the relative difficulty of the Maze and Melody contexts. We classified participants as Melody-first or Maze-first based on the first training context they encountered, then compared mean sequence RTs for Maze and Melody trials.

Across both start-order groups, Melody trials were slower than Maze trials. Participants who started with Melody showed mean RTs of 5645 ms in Melody and 4037 ms in Maze. Participants who started with Maze showed mean RTs of 5764 ms in Melody and 4866 ms in Maze. A repeated-measures ANOVA showed a reliable main effect of context, $F(1, 30) = 70.18$, $p < .001$, $\eta^2 = .09$, indicating that Melody was slower than Maze overall.

Start order also moderated the size of this context effect, $F(1, 30) = 5.80$, $p = .022$, $\eta^2 = .008$. Difference-score analyses showed the same pattern. Maze-minus-Melody scores were negative in both groups, confirming that Melody was slower than Maze regardless of start order. However, this difference was larger for participants who started with Melody than for participants who started with Maze (Melody-first: −1607 ms; Maze-first: −898 ms), $F(1, 30) = 5.80$, $p = .022$, $\eta^2 = .16$. Thus, starting order did not reverse which context was slower. Melody remained slower than Maze in both groups. Instead, the start order changed the magnitude of the context difference.

**Approach for Parameter Necessity Analyses for State-Space Model**

We used nested model comparisons to test which components of the shared-state model were necessary to explain the single-context human learning curves. This model contained one latent performance state and one latent fatigue state.

The state-space parameters were $A, \quad C, \quad \tau_{\text{rec}}$, where $A$ controlled the learning rate of the performance state, $C$ controlled fatigue accumulation within a trial, and $\tau_{\text{rec}}$ controlled fatigue recovery across breaks.

The null model fixed these parameters to values that removed learning and fatigue accumulation, $A = 1, C = 0, \tau_{\text{rec}} = 60$. We then fit nested variants of the model and compared them using BIC. Lower BIC indicates a better tradeoff between model fit and model complexity. The main test of parameter necessity was the leave-one-parameter-out ablation analysis. We started from the full lumped model,

$$\mathcal{M}_{\text{full}} = \{A, C, \tau_{\text{rec}}\}.$$

We then refit reduced models in which one parameter was fixed to its null value while the remaining state-space parameters and decoder coefficients were re-estimated

$$\mathcal{M}_{-A} = \{C, \tau_{\text{rec}}\},$$

$$\mathcal{M}_{-C} = \{A, \tau_{\text{rec}}\},$$

$$\mathcal{M}_{-\tau} = \{A, C\}.$$

We fit a stepwise sequence of nested models. This sequence started with the null model and then added $A$, $C$, and $\tau_{\text{rec}}$. The stepwise analysis shows how much each additional component improved the model as complexity was gradually increased. The ablation analysis provides a complementary test of necessity because the remaining parameters were allowed to compensate for the omitted parameter.

**State Space Parameter Necessity**

Across the three single-context conditions, learning was consistently necessary, whereas support for fatigue accumulation and recovery varied (Supplementary Table 1). In the Maze condition, the learning-only model was preferred by BIC (1267.63) over the full model (1274.99), although including fatigue and recovery modestly reduced RSS from 5,647,241 to 5,545,284. Neither model fully captured the within-block fluctuations observed during later training. By contrast, the full model substantially improved the Melody fit relative to learning alone (RSS = 3,155,435 vs. 4,499,002; ΔBIC = 22.93), while adding fatigue accumulation improved the No Context fit (RSS = 8,674,063 vs. 9,508,256; ΔBIC = 3.76). Thus, although the contribution of each component varied across conditions, fatigue accumulation and recovery were necessary to adequately describe performance in at least some conditions. We therefore retained the full learning–fatigue–recovery structure in all alternating-context models, ensuring that comparisons among Lump, Cross, and Split reflected differences in learning-state architecture rather than differences in the modeled fatigue process.

**Supplementary Table 1. Stepwise inclusion analysis for the lumped state model**

| Dataset | Model | BIC | ΔBIC |
|---|---|---|---|
| No Context | Null | 1391.63 | 80.88 |
| No Context | A | 1314.52 | 3.76 |
| No Context | **A + C** | **1310.75** | **0** |
| No Context | A + C + $\tau_{rec}$ | 1315.17 | 4.42 |
| Maze | Null | 1448.81 | 181.18 |
| Maze | **A** | **1267.63** | **0** |
| Maze | A + C | 1272.13 | 4.50 |
| Maze | A + C + $\tau_{rec}$ | 1274.99 | 7.36 |
| Melody | Null | 1464.85 | 240.61 |
| Melody | A | 1247.17 | 22.93 |
| Melody | A + C | 1230.44 | 6.20 |
| Melody | **A + C + $\tau_{rec}$** | **1224.24** | **0** |

For the stepwise analysis, ΔBIC is computed relative to the best stepwise model within each dataset.

**Supplementary Table 2. All BICs for State-Space Model Fits**

| Data | State Space Model | BIC |
|---|---|---|
| Maze–Melody | Lump | 2832 |
| Maze–Melody | Cross | 2719 |
| **Maze–Melody** | **Split** | **2708** |
| High-Scale | Lump | -400 |
| **High-Scale** | **Cross** | **-858** |
| High-Scale | Split | -819 |
| **Low-Scale** | **Lump** | **-1211** |
| Low-Scale | Cross | -1085 |
| Low-Scale | Split | -1049 |
| Low-Scale + High Context | Lump | -1133 |
| **Low-Scale + High Context** | **Cross** | **-1518** |
| Low-Scale + High Context | Split | -1151 |
| **High-Scale + Low Context** | **Lump** | **-912** |
| High-Scale + Low Context | Cross | -839 |
| High-Scale + Low Context | Split | -804 |

**Supplementary Table 3. State-space parameter values**

| Group | Counterbalance | Model | Parameter | Estimate |
|---|---|---|---|---|
| No Context | — | Single-state | A | 0.981 [0.971, 0.999] |
| No Context | — | Single-state | C | 493 [102, 2409] |
| No Context | — | Single-state | $\tau_{rec}$ | 44 [0, 2751] |

| | | | | |
|---|---|---|---|---|
| Maze | — | Single-state | A | 0.943 [0.935, 0.964] |
| Maze | — | Single-state | C | 118 [0, 356] |
| Maze | — | Single-state | $\tau_{rec}$ | 0.30 [0, 2212] |
| Melody | — | Single-state | A | 0.948 [0.942, 0.962] |
| Melody | — | Single-state | C | 429 [309, 634] |
| Melody | — | Single-state | $\tau_{rec}$ | 0 [0, 42] |
| Maze-Melody | Maze-first | Lump | A | 0.976 [0.954, 0.994] |
| Maze-Melody | Maze-first | Lump | C | 247 [0, 1820] |
| Maze-Melody | Maze-first | Lump | $\tau_{rec}$ | 134 [0, 3600] |
| Maze-Melody | Maze-first | Cross | $A_{Maze}$ | 0.920 [0.874, 0.934] |
| Maze-Melody | Maze-first | Cross | $A_{Cross}$ | 0.000 [0.000, 0.031] |
| Maze-Melody | Maze-first | Cross | $A_{Melody}$ | 0.959 [0.946, 0.965] |
| Maze-Melody | Maze-first | Cross | C | 376 [182, 622] |
| Maze-Melody | Maze-first | Cross | $\tau_{rec}$ | 0 [0, 25] |
| Maze-Melody | Maze-first | Split | $A_{Maze}$ | 0.920 [0.904, 0.936] |
| Maze-Melody | Maze-first | Split | $A_{Melody}$ | 0.958 [0.950, 0.966] |
| Maze-Melody | Maze-first | Split | C | 377 [158, 631] |
| Maze-Melody | Maze-first | Split | $\tau_{rec}$ | 0 [0, 21] |
| Maze-Melody | Melody-first | Lump | A | 0.948 [0.937, 0.973] |
| Maze-Melody | Melody-first | Lump | C | 106 [0, 663] |
| Maze-Melody | Melody-first | Lump | $\tau_{rec}$ | 25 [0, 3097] |
| Maze-Melody | Melody-first | Cross | $A_{Maze}$ | 0.872 [0.834, 0.899] |
| Maze-Melody | Melody-first | Cross | $A_{Cross}$ | 0.000 [0.000, 0.034] |
| Maze-Melody | Melody-first | Cross | $A_{Melody}$ | 0.956 [0.935, 0.962] |
| Maze-Melody | Melody-first | Cross | C | 445 [89, 867] |
| Maze-Melody | Melody-first | Cross | τrec | 0 [0, 53] |
| Maze-Melody | Melody-first | Split | $A_{Maze}$ | 0.872 [0.842, 0.901] |
| Maze-Melody | Melody-first | Split | $A_{Melody}$ | 0.956 [0.945, 0.964] |

| | | | | |
|---|---|---|---|---|
| Maze-Melody | Melody-first | Split | C | 445 [78, 859] |
| Maze-Melody | Melody-first | Split | $\tau_{rec}$ | 0 [0, 33] |
| High-Scale + Low Context | Maze-first | Cross | $A_{Maze}$ | 0.867 [0.738, 0.947] |
| High-Scale + Low Context | Maze-first | Cross | $A_{Melody}$ | 0.767 [0.600, 0.946] |
| High-Scale + Low Context | Maze-first | Cross | $A_{cross}$ | 0.069 [0.000, 0.423] |
| High-Scale + Low Context | Maze-first | Lumped | A | 0.936 [0.826, 0.999] |
| High-Scale + Low Context | Maze-first | Split | $A_{Maze}$ | 0.925 [0.850, 0.995] |
| High-Scale + Low Context | Maze-first | Split | $A_{Melody}$ | 0.833 [0.600, 0.992] |
| High-Scale + Low Context | Melody-first | Cross | $A_{Maze}$ | 0.685 [0.600, 0.944] |
| High-Scale + Low Context | Melody-first | Cross | $A_{Melody}$ | 0.756 [0.677, 0.945] |
| High-Scale + Low Context | Melody-first | Cross | $A_{cross}$ | 0.194 [0.053, 0.279] |
| High-Scale + Low Context | Melody-first | Lumped | A | 0.969 [0.920, 0.999] |
| High-Scale + Low Context | Melody-first | Split | $A_{Maze}$ | 0.922 [0.756, 0.988] |
| High-Scale + Low Context | Melody-first | Split | $A_{Melody}$ | 0.953 [0.899, 0.990] |
| High-Scale | Maze-first | Cross | $A_{Maze}$ | 0.948 [0.831, 0.985] |
| High-Scale | Maze-first | Cross | $A_{Melody}$ | 0.818 [0.600, 0.936] |
| High-Scale | Maze-first | Cross | $A_{cross}$ | 0.047 [0.000, 0.148] |
| High-Scale | Maze-first | Lumped | A | 0.976 [0.949, 0.999] |
| High-Scale | Maze-first | Split | $A_{Maze}$ | 0.975 [0.943, 0.997] |
| High-Scale | Maze-first | Split | $A_{Melody}$ | 0.856 [0.600, 0.942] |
| High-Scale | Melody-first | Cross | $A_{Maze}$ | 0.879 [0.651, 0.952] |
| High-Scale | Melody-first | Cross | $A_{Melody}$ | 0.956 [0.867, 0.987] |
| High-Scale | Melody-first | Cross | $A_{cross}$ | 0.028 [0.000, 0.098] |
| High-Scale | Melody-first | Lumped | A | 0.980 [0.951, 0.999] |
| High-Scale | Melody-first | Split | $A_{Maze}$ | 0.900 [0.694, 0.961] |
| High-Scale | Melody-first | Split | $A_{Melody}$ | 0.973 [0.942, 0.993] |
| Low-Scale + High Context | Maze-first | Cross | $A_{Maze}$ | 0.999 [0.994, 0.999] |
| Low-Scale + High Context | Maze-first | Cross | $A_{Melody}$ | 0.995 [0.988, 0.999] |
| Low-Scale + High Context | Maze-first | Cross | $A_{cross}$ | 0.064 [0.058, 0.071] |
| Low-Scale + High Context | Maze-first | Lumped | A | 0.999 [0.999, 0.999] |
| Low-Scale + High Context | Maze-first | Split | $A_{Maze}$ | 0.997 [0.994, 0.999] |
| Low-Scale + High Context | Maze-first | Split | $A_{Melody}$ | 0.996 [0.992, 0.999] |
| Low-Scale + High Context | Melody-first | Cross | $A_{Maze}$ | 0.999 [0.998, 0.999] |
| Low-Scale + High Context | Melody-first | Cross | $A_{Melody}$ | 0.994 [0.984, 0.999] |
| Low-Scale + High Context | Melody-first | Cross | $A_{cross}$ | 0.055 [0.048, 0.062] |
| Low-Scale + High Context | Melody-first | Lumped | A | 0.999 [0.999, 0.999] |
| Low-Scale + High Context | Melody-first | Split | $A_{Maze}$ | 0.998 [0.992, 0.999] |
| Low-Scale + High Context | Melody-first | Split | $A_{Melody}$ | 0.998 [0.994, 0.999] |
| Low-Scale | Maze-first | Cross | $A_{Maze}$ | 0.863 [0.861, 0.866] |
| Low-Scale | Maze-first | Cross | $A_{Melody}$ | 0.859 [0.857, 0.863] |
| Low-Scale | Maze-first | Cross | $A_{cross}$ | 0.110 [0.107, 0.112] |
| Low-Scale | Maze-first | Lumped | A | 0.987 [0.986, 0.989] |
| Low-Scale | Maze-first | Split | $A_{Maze}$ | 0.975 [0.973, 0.978] |
| Low-Scale | Maze-first | Split | $A_{Melody}$ | 0.970 [0.967, 0.973] |
| Low-Scale | Melody-first | Cross | $A_{Maze}$ | 0.860 [0.857, 0.864] |
| Low-Scale | Melody-first | Cross | $A_{Melody}$ | 0.863 [0.861, 0.867] |
| Low-Scale | Melody-first | Cross | $A_{cross}$ | 0.110 [0.108, 0.113] |
| Low-Scale | Melody-first | Lumped | A | 0.988 [0.987, 0.990] |
| Low-Scale | Melody-first | Split | $A_{Maze}$ | 0.972 [0.968, 0.975] |

| | | | | |
|---|---|---|---|---|
| Low-Scale | Melody-first | Split | $A_{Melody}$ | 0.976 [0.974, 0.979] |

*Note.* Values are bootstrap means with 95% percentile confidence intervals in brackets. Each interval is based on 500 bootstrap resamples.

**Supplementary Table 4 Parameter Recovery**

| Condition | Counterbalance | Model | Parameter | True | Recovered | RMSE |
|---|---|---|---|---|---|---|
| No Context | | Lumped | A | 0.986922 | 0.986921 | 0.000001 |
| No Context | | Lumped | C | 280.72313 | 280.724786 | 0.001656 |
| No Context | | Lumped | $\tau_{rec}$ | 0 | 0 | 0 |
| Maze | — | Lumped | A | 0.943313 | 0.943313 | 0.0000002 |
| Maze | — | Lumped | C | 118.40123 | 118.398199 | 0.003037 |
| Maze | — | Lumped | $\tau_{rec}$ | 0 | 0 | 0 |
| Melody | — | Lumped | A | 0.908 | 0.881 | 0.027 |
| Melody | — | Lumped | C | 429.46433 | 429.470942 | 0.006610 |
| Melody | — | Lumped | $\tau_{rec}$ | 0 | 0 | 0 |
| Maze-Melody | Maze-first | Lumped | A | 0.9762 | 0.9715 | 0.0122 |
| Maze-Melody | Maze-first | Lumped | C | 247.5 | 341.8 | 475.2 |
| Maze-Melody | Maze-first | Lumped | $\tau_{rec}$ | 134.3 | 623.0 | 1258.2 |
| Maze-Melody | Maze-first | Cross | $A_{cross}$ | 0.9200 | 0.9145 | 0.0144 |
| Maze-Melody | Maze-first | Cross | $A_{diag}$ | 0.0000 | 0.0048 | 0.0097 |
| Maze-Melody | Maze-first | Cross | $C_{Maze}$ | 0.9585 | 0.9566 | 0.0048 |
| Maze-Melody | Maze-first | Cross | $C_{Melody}$ | 376.3 | 388.1 | 111.9 |
| Maze-Melody | Maze-first | Cross | $\tau_{rec}$ | 0.0 | 5.8 | 57.0 |
| Maze-Melody | Maze-first | Split | $A_{diag}$ | 0.9199 | 0.9211 | 0.0100 |
| Maze-Melody | Maze-first | Split | $C_{Maze}$ | 0.9585 | 0.9584 | 0.0047 |
| Maze-Melody | Maze-first | Split | $C_{Melody}$ | 376.6 | 396.6 | 122.7 |
| Maze-Melody | Maze-first | Split | $\tau_{rec}$ | 0.0 | 3.8 | 7.3 |
| Maze-Melody | Melody-first | Lumped | A | 0.9482 | 0.9518 | 0.0110 |

| | | | | | | |
|---|---|---|---|---|---|---|
| Maze-Melody | Melody-first | Lumped | C | 105.8 | 203.1 | 243.5 |
| Maze-Melody | Melody-first | Lumped | $\tau_{rec}$ | 25.2 | 368.8 | 928.8 |
| Maze-Melody | Melody-first | Cross | $A_{cross}$ | 0.8723 | 0.8658 | 0.0201 |
| Maze-Melody | Melody-first | Cross | $A_{diag}$ | 0.0000 | 0.0058 | 0.0107 |
| Maze-Melody | Melody-first | Cross | $C_{Maze}$ | 0.9556 | 0.9509 | 0.0087 |
| Maze-Melody | Melody-first | Cross | $C_{Melody}$ | 444.6 | 504.6 | 180.9 |
| Maze-Melody | Melody-first | Cross | $\tau_{rec}$ | 0.0 | 24.4 | 211.1 |
| Maze-Melody | Melody-first | Split | $A_{diag}$ | 0.8723 | 0.8708 | 0.0177 |
| Maze-Melody | Melody-first | Split | $C_{Maze}$ | 0.9556 | 0.9544 | 0.0049 |
| Maze-Melody | Melody-first | Split | $C_{Melody}$ | 444.6 | 489.7 | 184.2 |
| Maze-Melody | Melody-first | Split | $\tau_{rec}$ | 0.0 | 13.2 | 161.6 |
| High-Scale | Maze-first | Lumped | A | 0.977137 | 0.977133 | 0.000003 |
| High-Scale | Maze-first | Cross | $A_{Maze}$ | 0.966003 | 0.965980 | 0.000023 |
| High-Scale | Maze-first | Cross | $A_{Cross}$ | 0.031361 | 0.031406 | 0.000045 |
| High-Scale | Maze-first | Cross | $A_{Melody}$ | 0.857347 | 0.857367 | 0.000020 |
| High-Scale | Maze-first | Split | $A_{Maze}$ | 0.978863 | 0.978857 | 0.000006 |
| High-Scale | Maze-first | Split | $A_{Melody}$ | 0.879180 | 0.879133 | 0.000047 |
| High-Scale | Melody-first | Lumped | A | 0.980294 | 0.980291 | 0.000003 |
| High-Scale | Melody-first | Cross | $A_{Maze}$ | 0.903813 | 0.903691 | 0.000121 |
| High-Scale | Melody-first | Cross | $A_{Cross}$ | 0.015702 | 0.015806 | 0.000104 |
| High-Scale | Melody-first | Cross | $A_{Melody}$ | 0.968292 | 0.968222 | 0.000070 |
| High-Scale | Melody-first | Split | $A_{Maze}$ | 0.913759 | 0.913691 | 0.000067 |
| High-Scale | Melody-first | Split | $A_{Melody}$ | 0.975735 | 0.975715 | 0.000020 |
| Low-Scale | Maze-first | Lumped | A | 0.987467 | 0.987468 | 0.000000 |
| Low-Scale | Maze-first | Cross | $A_{Maze}$ | 0.863259 | 0.861252 | 0.002008 |

| | | | | | | |
|---|---|---|---|---|---|---|
| Low-Scale | Maze-first | Cross | $A_{Cross}$ | 0.109257 | 0.111167 | 0.001910 |
| Low-Scale | Maze-first | Cross | $A_{Melody}$ | 0.859651 | 0.857676 | 0.001975 |
| Low-Scale | Maze-first | Split | $A_{Maze}$ | 0.974944 | 0.974229 | 0.000715 |
| Low-Scale | Maze-first | Split | $A_{Melody}$ | 0.969758 | 0.968854 | 0.000903 |
| Low-Scale | Melody-first | Lumped | A | 0.988195 | 0.988194 | 0.000001 |
| Low-Scale | Melody-first | Cross | $A_{Maze}$ | 0.858474 | 0.853969 | 0.004505 |
| Low-Scale | Melody-first | Cross | $A_{Cross}$ | 0.111938 | 0.116298 | 0.004359 |
| Low-Scale | Melody-first | Cross | $A_{Melody}$ | 0.861709 | 0.857130 | 0.004578 |
| Low-Scale | Melody-first | Split | $A_{Maze}$ | 0.972035 | 0.970980 | 0.001055 |
| Low-Scale | Melody-first | Split | $A_{Melody}$ | 0.976706 | 0.975852 | 0.000853 |
| Low-Scale + High Context | Maze-first | Lumped | A | 0.999000 | 0.999000 | 0.000000 |
| Low-Scale + High Context | Maze-first | Cross | $A_{Maze}$ | 0.999000 | 0.999000 | 0.000000 |
| Low-Scale + High Context | Maze-first | Cross | $A_{Cross}$ | 0.062775 | 0.062774 | 0.000001 |
| Low-Scale + High Context | Maze-first | Cross | $A_{Melody}$ | 0.995523 | 0.995526 | 0.000003 |
| Low-Scale + High Context | Maze-first | Split | $A_{Maze}$ | 0.998357 | 0.989836 | 0.008521 |
| Low-Scale + High Context | Maze-first | Split | $A_{Melody}$ | 0.997960 | 0.986994 | 0.010967 |
| Low-Scale + High Context | Melody-first | Lumped | A | 0.999000 | 0.999000 | 0.000000 |
| Low-Scale + High Context | Melody-first | Cross | $A_{Maze}$ | 0.999000 | 0.999000 | 0.000000 |
| Low-Scale + High Context | Melody-first | Cross | $A_{Cross}$ | 0.053619 | 0.053620 | 0.000001 |
| Low-Scale + High Context | Melody-first | Cross | $A_{Melody}$ | 0.994343 | 0.994348 | 0.000005 |

| | | | | | | |
|---|---|---|---|---|---|---|
| Low-Scale + High Context | Melody-first | Split | $A_{Maze}$ | 0.998724 | 0.987246 | 0.011477 |
| Low-Scale + High Context | Melody-first | Split | $A_{Melody}$ | 0.999000 | 0.990288 | 0.008712 |
| High-Scale + Low Context | Maze-first | Lumped | A | 0.939645 | 0.939640 | 0.000005 |
| High-Scale + Low Context | Maze-first | Cross | $A_{Maze}$ | 0.865598 | 0.865550 | 0.000048 |
| High-Scale + Low Context | Maze-first | Cross | $A_{Cross}$ | 0.052601 | 0.052634 | 0.000033 |
| High-Scale + Low Context | Maze-first | Cross | $A_{Melody}$ | 0.780382 | 0.780333 | 0.000049 |
| High-Scale + Low Context | Maze-first | Split | $A_{Maze}$ | 0.919218 | 0.919214 | 0.000004 |
| High-Scale + Low Context | Maze-first | Split | $A_{Melody}$ | 0.838170 | 0.838163 | 0.000007 |
| High-Scale + Low Context | Melody-first | Lumped | A | 0.971826 | 0.971822 | 0.000004 |
| High-Scale + Low Context | Melody-first | Cross | $A_{Maze}$ | 0.605023 | 0.604901 | 0.000123 |
| High-Scale + Low Context | Melody-first | Cross | $A_{Cross}$ | 0.265202 | 0.265299 | 0.000097 |
| High-Scale + Low Context | Melody-first | Cross | $A_{Melody}$ | 0.686549 | 0.686441 | 0.000108 |
| High-Scale + Low Context | Melody-first | Split | $A_{Maze}$ | 0.937486 | 0.937367 | 0.000119 |
| High-Scale + Low Context | Melody-first | Split | $A_{Melody}$ | 0.955826 | 0.955747 | 0.000079 |

*Note.* True values are the fitted parameter values used to generate simulated data. Recovered values are the estimates obtained after refitting the model to the simulated data. Error = |Recovered - True|.
